# CXCR2 expression during melanoma tumorigenesis controls transcriptional programs that facilitate tumor growth

**DOI:** 10.1101/2023.02.22.529548

**Authors:** J Yang, K Bergdorf, C Yan, W Luo, SC Chen, D Ayers, Q Liu, X Liu, M Boothby, SM Groves, AN Oleskie, X Zhang, DY Maeda, JA Zebala, V Quaranta, A Richmond

## Abstract

**Background:** Though the CXCR2 chemokine receptor is known to play a key role in cancer growth and response to therapy, a direct link between expression of CXCR2 in tumor progenitor cells during induction of tumorigenesis has not been established.

**Methods:** To characterize the role of CXCR2 during melanoma tumorigenesis, we generated tamoxifen-inducible tyrosinase-promoter driven *Braf^V600E^/Pten^-/-^/Cxcr2^-/-^* and *NRas^Q61R^/INK4a^-/-^/Cxcr2^-/-^* melanoma models. In addition, the effects of a CXCR1/CXCR2 antagonist, SX-682, on melanoma tumorigenesis were evaluated in *Braf^V600E^/Pten^-/-^* and *NRas^Q61R^/INK4a^-/-^* mice and in melanoma cell lines. Potential mechanisms by which *Cxcr2* affects melanoma tumorigenesis in these murine models were explored using RNAseq, mMCP-counter, ChIPseq, and qRT-PCR; flow cytometry, and reverse phosphoprotein analysis (RPPA).

**Results:** Genetic loss of *Cxcr2* or pharmacological inhibition of CXCR1/CXCR2 during melanoma tumor induction resulted in key changes in gene expression that reduced tumor incidence/growth and increased anti-tumor immunity. Interestingly, after *Cxcr2* ablation, *Tfcp2l1*, a key tumor suppressive transcription factor, was the only gene significantly induced with a log_2_ fold-change greater than 2 in these three different melanoma models.

**Conclusions:** Here, we provide novel mechanistic insight revealing how loss of *Cxcr2* expression/activity in melanoma tumor progenitor cells results in reduced tumor burden and creation of an anti-tumor immune microenvironment. This mechanism entails an increase in expression of the tumor suppressive transcription factor, *Tfcp2l1,* along with alteration in the expression of genes involved in growth regulation, tumor suppression, stemness, differentiation, and immune modulation. These gene expression changes are coincident with reduction in the activation of key growth regulatory pathways, including AKT and mTOR.

## Introduction

Chemokines and their receptors have been shown to play an essential role in regulating tumor growth, progression, metastasis, and response to immunotherapy (1, 2, 3, 4). Though chemokines were initially identified as chemoattractants used to guide leukocyte migration, there has been increasing evidence that they can regulate other functions in a broader array of cell types, including cancer cells (5).

The CXCR1/CXCR2 ligand-receptor axis has been widely characterized as a driver of aggressive behavior in many cancer types, including breast, prostate, melanoma, lung, colorectal, pancreatic, and renal cancers(6). CXCR1/CXCR2 ligands, including CXCL1-3, 5-8 are produced by endothelial cells, tumor-associated macrophages, cancer-associated fibroblasts, adipocytes, and cancer cells(6). These CXCR1 and CXCR2 ligands play a significant role in the recruitment of neutrophils and myeloid-derived suppressor cells (MDSCs) to the tumor microenvironment (TME), both of which are associated with poor outcomes(7, 8, 9). In addition to altering the tumor immune microenvironment, these chemokine ligands can also activate phosphatidylinositol-3-βkinase (PI3K), phospholipase-Cβ, calcium mobilization, mitogen-activated protein kinase (MAPK), protein kinase B (AKT), transcription factors like NF-κ expression on tumor cells. These chemokine responses have been linked to tumor cell survival, proliferation, migration, as well as angiogenesis(6, 10, 11).

Many cancer cells exhibit induction or increased expression of multiple ligands for both CXCR1 (CXCL1-3, 5-8) and CXCR2 (CXCL1-3, 5 and 7). Moreover, CXCR1 and CXCR2 are differentially expressed in human tissues, though in mouse, CXCR2 is the predominant receptor mediating response to the murine chemokine ligands during inflammation, angiogenesis, and tumor growth (CXCL1,2,3 and 5, also known as KC, MIP2α, MIP2β, and LIX)(12, 13). In addition to a function in the attraction of hematopoietic cells that influence the tumor microenvironment and tumor progression, it has been suggested that these receptors may exert autocrine effects on tumor growth. In the case of melanoma, mouse xenograft models provide compelling evidence that tumor cells take advantage of CXCR2 ligand expression to either suppress the anti-tumor immune response or to induce tumor growth and angiogenesis, alter the TME, and facilitate metastasis (3, 14)

The CXCR1/CXCR2 signaling nexus directly influences the sensitivity of tumor cells to chemotherapies by altering pathways associated with apoptosis and multidrug resistance (15, 16), resulting in a poor prognosis in human cancer studies (17, 18). The past decade has witnessed the generation and development of antagonists to CXCR1 and CXCR2, and multiple clinical trials are underway investigating the therapeutic potential of targeting this signaling axis in inflammatory disorders and cancers (NCT03161431, NCT04245397, NCT03400332) (19, 20, 21, 22).

We previously demonstrated that targeted deletion of *Cxcr2* in myeloid cells or systemic treatment with the CXCR1/CXCR2 antagonist SX-682 conferred anti-tumor immunity via reduction of MDSC infiltration into the TME and enhanced CD8+ T cell activation (9). However, it remains controversial as to whether there is a direct function of either or both CXCR1 and CXCR2 on the growth of the cancer cells, and if so, which of these receptors are involved and what mechanisms are employed. To clarify the concept of an autocrine role for CXCR2 and its ligands in melanoma progenitor cells, we used inducible, autochthonous models of malignant melanoma in mice. Using two distinct modes of triggering the formation of malignant melanoma (23, 24) (25), we found that tumor onset, growth, and outcome accompanied changes in the tumor microenvironment and gene expression when *Cxcr2* was deleted in melanoma precursor cells. Similar results were identified when Cxcr1/Cxcr2 were inhibited with SX-682 during tumorigenesis. Remarkably, an analysis of common gene expression changes due to loss or inhibition of Cxcr2 during tumorigenesis converged on one, but only one, gene -- the tumor suppressive transcription factor *Tfcp2l1*. These data indicate that a major mechanism by which Cxcr2 inhibition regulates melanoma tumor growth is via induction of a key transcription factor with tumor suppressive activity, *Tfcp2l1*.

## Methods

### Establishment of inducible melanoma mouse models

All procedures involving animals were approved by the Vanderbilt University Institutional Animal Care and Use Committee (IACUC). We utilized the inducible *Braf^V600E^/PTEN^-/-^* melanoma model in C57BL/6 mice (23), where the underlying genetic background includes *Tyr-Cre^ER+^:: Braf^CA^::Pten^lox4^*^-5/Lox4-5^. CXCR2^f/f^ mice *(C57BL/6-CXCR2^tm1RMra/J^)* were obtained from Jackson Laboratories (#024638) and bred to mT/mG mice (#007907, Jackson Laboratories), which harbor a two-color fluorescent Cre-reporter allele to enable GFP-based tumor imaging (Figure S2A,C) (26). In crossing the *Braf^V600E^/PTEN^-/-^* mice with CXCR2^fl/fl^ mT/mG mice, Tyr-*Cre^ER+^::Braf^CA^::Pten^lox4-5/Lox4-5^::mT/mG::Cxcr2^fl/fl^* mice and *Cre^ER+^:: Braf^CA^::Pten^lox4-5/Lox4-5^::mT/mG::Cxcr2^WT^* mice were generated. Upon administration of 4-HT (#6278, Sigma), Cre-recombinase expression is induced in tyrosinase (*Tyr*) expressing cells, leading to expression of the *Braf^V600E^* transgene and deletion of exons 4 and 5 of *Pten* specifically in tyrosinase expressing melanocytes (Figure S2B)(23). Palpable tumors arise within one month post 4-HT induction (Figure S2B, C). Tyr-Cre targeting of melanocytes in hair follicles was verified by H&E staining and GFP expression (Figure S2D).

To generate an inducible *NRas* mutant/*Ink4a* deletion/CXCR2 knockout melanoma mouse model, we utilized the *TpN^61R^* model from Burd et al., which recapitulates the genetics of *NRas^Q61R^/INK4a^-/-^* mutant human melanoma and demonstrates sensitivity to UV-induced melanoma (25). In this model, expression of mutant *NRas* and loss of *Ink4a* are under the control of the *Tyr*-promoter enhancer *(Tyr-Cre^ER^::NRas^Q61R^::Ink4a^-/-^).* These mice were crossed with C57BL/6 *Cxcr2^f/f^* mice. Heterozygous offspring were crossed to generate *Tyr-Cre^ER^::NRas^Q61R^::Ink4a^fl/fl^::Cxcr^fl/fl^ and Tyr-Cre^ER^::NRas^Q61R^::Ink4a^fl/fl^::Cxcr2^WT^* littermates. Newborn mice (1-2 days of age) receive one topical administration of 2μl of 20mM 4HT on the back followed by exposure to 4.5K J/m2 UVB radiation (312NM 2X8 Watt tubes& Filter, Cat. # EB-280C) on day three. Tumor development was followed for 5 months. **All other standard methods are in the Supplemental Materials.**

## Results

### CXCR2 Correlates with Poor Prognosis in Patient Populations and Response to Checkpoint inhibitors

Using the available Gene Expression Omnibus (GEO) cohort, we evaluated *CXCR1, CXCR2,* and *CXCL1-3, 5* and *8* (CXCR1/CXCR2 ligands) expression in nevi and melanoma. *CXCR1* and *CXCR2* mRNA exhibited a trend toward increased expression in melanoma compared to nevi, but these differences were not statistically significant (Figure 1A). However, C*XCL1*, *CXCL2*, *CXCL3*, *CXCL5* and *CXCL8* mRNAs were significantly upregulated in melanoma samples compared to benign nevi (Figure 1B). Furthermore, there were no significant differences in *CXCR1* and *CXCR2* expression among nevi and melanoma tumors when stratified by *BRAF* or *NRAS* mutation status (Figure S1A, B). However, since the number of samples available for analysis of mutation status was small, these findings should be interpreted cautiously.

**Fig 1.**
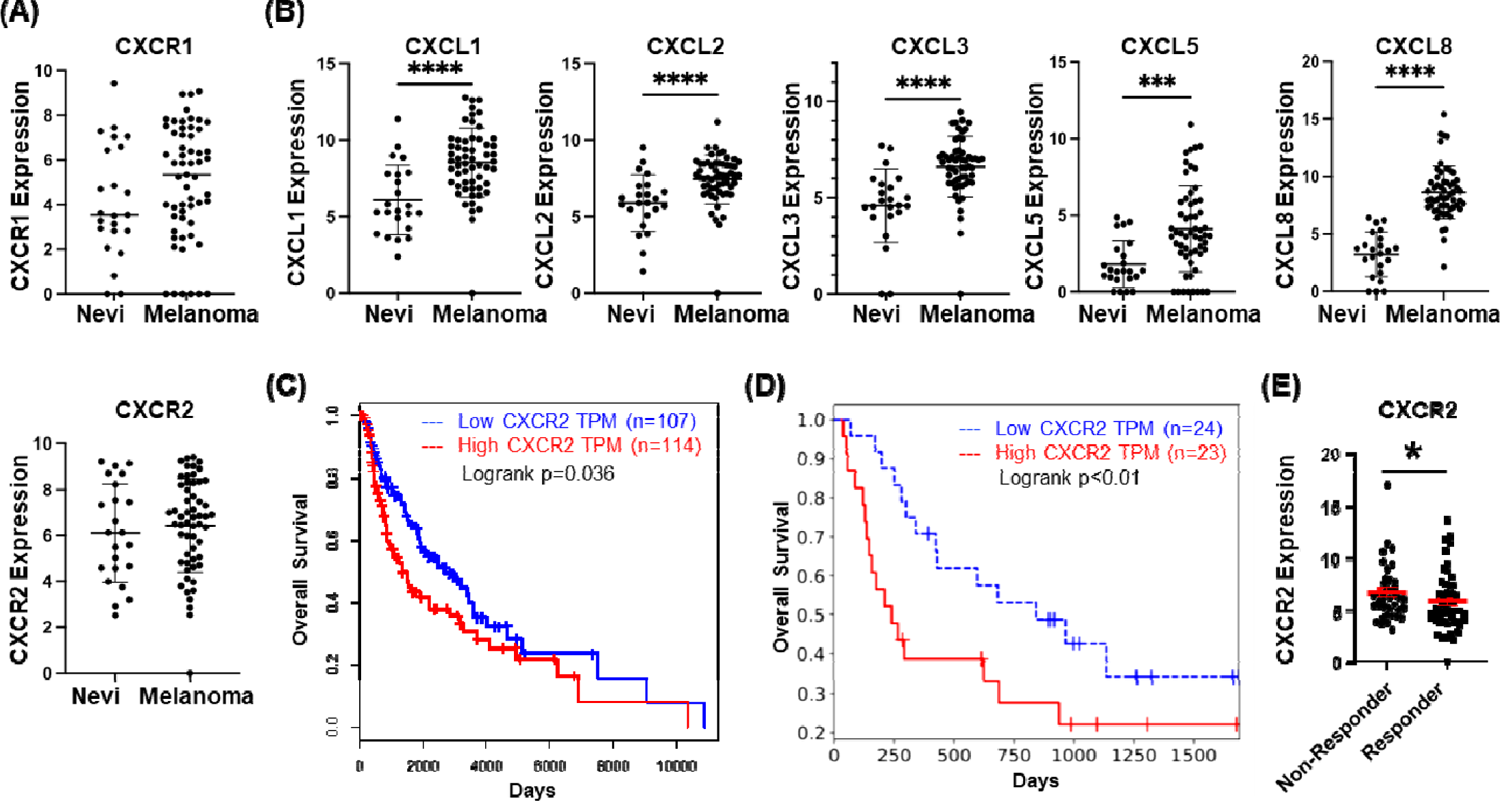
CXCR2 is associated with tumorigenesis and poor prognosis. **a** GEO dataset analysis of expression of CXCR1 and CXCR2 in nevi as compared to melanoma lesions (not significant, Welch’s t-test). **b** GEO dataset analysis of expression of CXCL1, CXCL2, CXCL3, CXCL5 and CXCL8 in nev and melanoma tissues (significance determined by Welch’s t-test). **c** Overall survival plot of melanoma patients from the TCGA SKCM dataset indicates significantly improved survival (p=0.035, log-rank test) in the lowest quartile of CXCR2 expression (blue, n=107) compared to the highest quartile (red, n=114). **d** Analysis of survival of 25 melanoma patients treated with anti-PD1 in relation to high (red) or low (blue) expression of CXCR2 [p<0.01, log-rank test; (27)].. **e** Re-analysis of the Riaz RNA-seq database shows CXCR2 expression is lower in melanoma patients who responded to anti-PD1 treatment (p<0.05, Welch’s t-test).

CXCR2 has been associated with increased tumor growth and poor prognosis across multiple cancers(6). To define the relationship between *CXCR2* expression and the clinical prognosis of melanoma patients, we examined clinical data from the Cancer Genome Atlas (TCGA), and the skin cutaneous melanoma (SKCM) dataset using Gene Expression Profiling Interactive Analysis (GEPIA). Survival analysis comparing patients with high *CXCR2* expression (n=114) to patients with lower *CXCR2* expression (n=107) indicates that *CXCR2* expression correlates with decreased overall survival of melanoma patients (p=0.035, Figure 1C). Evaluation of survival in a patient cohort treated with anti-PD-1 therapy also suggests that patients with high *CXCR2* expression (n=24) exhibited poor prognosis in response to anti-PD-1 when compared with patients with low *CXCR2* expression (n=23, p<0.01; Figure 1D) (27). Finally, analysis of another immune checkpoint inhibitor-treated cohort showed that responding patients had significantly lower *CXCR2* expression than non-responders (Figure 1E, (p<0.05) (28). These data indicate that *CXCR2* expression correlates with poor therapeutic response in melanoma patients.

### CXCR2 Influences Tumor Differentiation Status and Enhances Tumor Growth

To evaluate the role of CXCR2 in *Braf^V600E^/Pten^-/-^* melanoma tumorigenesis, we crossed C57BL/6 *Tyr-CreER+::Braf^V600E^/Pten^fl/fl^::mT/mG*:: mice (*Braf^V600E^/Pten^-/-^)* (*23*) with C57BL/6 mice carrying a *Cxcr2^fl/fl^* allele (24) to produce *Tyr-CreER+::Braf^V600E^/Pten^fl/fl^::mT/mG::Cxcr2^-/-^* and *Tyr-CreER+::Braf^V600E^/Pten^fl/fl^::mT/mG::Cxcr2^WT^* littermates. Four-week-old mice were treated with 4-OH tamoxifen (4HT) to induce the tyrosinase promoter-driven Cre-recombinase. The resulting melanoma tumors that developed over 36 days were counted and measured. We observed that tumor burden and incidence (Figure 2A-C) were significantly reduced in *Braf^V600E^/Pten^-/-^* mice with melanocyte targeted deletion of *Cxcr2* (*Braf/Pten/Cxcr2^-/-^)* (271±361mm^3^, n=21) in comparison to *Braf^V600E^/Pten^-/-^* mice expressing CXCR2 *(Braf/Pten/Cxcr2^WT^)* (615±609mm^3^, n=24, p<0.05). The tumor number per mouse was also reduced upon melanocytic *Cxcr2* deletion (0.7±0.9 vs. 2.1±2.3, p<0.05). These data indicate that the Cxcr2 signal transduction pathway plays a role in the induction and growth of *Braf^V600E^/Pten^-/-^* melanoma.

**Fig 2.**
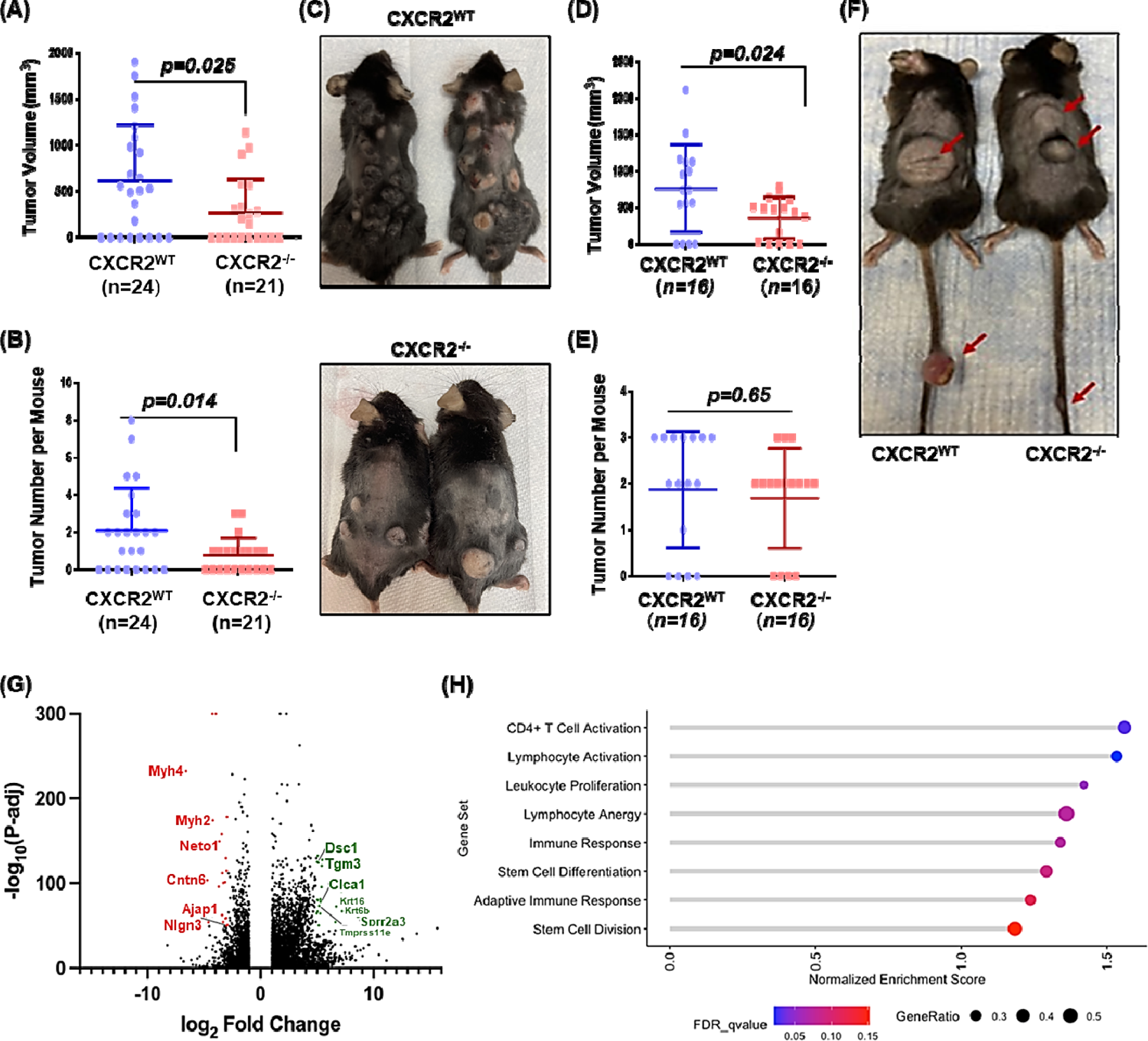
CXCR2 knockout decreases melanoma tumor burden. *Tyr-Cre^ER+^:: Braf^CA/+^::Pten ^ox4-5/lox4-5^::mT/mG* C57BL/6 mice were crossed with floxed *Cxcr2* mice to obtain mice with inducible tumors with or without CXCR2 expression Thirty-six days after 4-HT administration, **a** skin tumour volume and **b** count were recorded, and **c** mice were photographed (significance determined by Welch’s t-test). Similarly, *Tyr-Cre^ER+^::NRas^Q61R^::Ink4a^-/-^::mT/mG mG* mice were crossed with floxed *Cxcr2* mice, and resulting pups were treated with 4-HT on days 1 and 2 prior to UV irradiation on day 3 to initiate tumor formation (n=16/genotype). **d** Tumors were measured, **e** counted, and **f** mice were photographed (significance determined by Welch’s t-test). RNA was extracted from *Braf^V600E^/Pten^-/-^/Cxcr2^-/-^* and *Braf^V600E^/Pten^-/-^/Cxcr2^WT^* tumors and subjected to RNAseq analysis. **g** A volcano plot showing fold change and signific nce of differential gene expression in *Cxcr2^-/-^* tumors compared to *Cxcr2^WT^* tumors. **h** Gene set enrichment nalysis (GSEA) of RNAseq data identifies 8 gene sets enriched in *Cxcr2^-/-^* tumors. Point size indicates the gene ratio (percent of genes from the gene set contributing to the enrichment score) and point color represents the FDR q-value.

To determine whether *Cxcr2* is also important in *Nras/Ink4a* melanoma tumors, we crossed *Tyr-CreER+::NRas^Q61R^/Ink4a^-/-^::mT/mG::* mice (34) with the *Cxcr2^fl/fl^* mice (24). New-born pups (1-2 days old) were exposed to 4-HT, followed by ultraviolet (UV) irradiation on day three, and tumor growth was evaluated over five months. We observed significantly reduced tumor volume with deletion of *Cxcr2* (360±285mm^3^) when compared to *NRas/Ink4a Cxcr2^WT^* mice (764±601mm^3^) (Figure 2D-F, p<0.05, n=16). However, in contrast to the *Braf^V600E^/Pten^-/-^* model, the number of tumors per mouse was not significantly different between *Cxcr2^-/-^* (1.69±1.08) and *Cxcr2^WT^* mice (1.88±1.26, p=0.654). As the *NRas* GEM model requires UV irradiation in addition the genetic alterations, it is possible that UV irradiation induces additional oncogenic pathways that function independent of Cxcr2.

To elucidate the mechanism by which Cxcr2 perturbation in melanocytes could alter the initiation and growth of *Braf^V600E^/Pten^-/-^ (Braf/Pten)* melanoma, we examined the transcriptome of tumors arising in *Braf/Pten/Cxcr2^WT^* (n=7) and *Braf/Pten/Cxcr2*^-/-^ (n=8) mice via RNA sequencing (RNAseq) analysis (Figures 2G, S3A, S4). Interestingly, gene set enrichment analysis revealed that loss of *Cxcr2* expression in *Braf/Pten* tumors resulted in a significant increase in expression of genes involved in CD4+ T cell activation and lymphocyte activation, with a trend toward increased leukocyte proliferation, immune response, and stem cell differentiation (Figure 2H). However, there is also a paradoxical change in genes involved in lymphocyte anergy. Several genes were upregulated in *Braf/Pten/Cxcr2^-/-^* tumors, including those that are immune related, associated with stem cell differentiation, and those involved in tumor suppression. This gene analysis also revealed downregulation of genes involved in growth, proliferation, and cell cycle; immune-related genes; motility and cell adhesion; differentiation/stemness and tumor suppression (Figure S3A). Together, these RNA sequencing data imply that *Cxcr2^-/-^* tumors may become less cohesive/dense, with diminished invasive potential and growth signaling as modeled by Eikenberry *et al.* (*29*).

We next evaluated the RNAseq data from *Braf/Pten* mice with or without loss of CXCR2 expression in melanocytes and identified the most differentially expressed genes that are associated with favorable or unfavorable outcome in melanoma patients. We identified the top twenty growth related genes with reduced expression and the top twenty genes associated with inhibition of tumor growth and favorable outcome based on their log_2_ fold change and -log_10_ p-value (Figure S4). Key growth stimulatory (Figure 4A) and tumor suppressive genes (Figure 4B) are indicated by red arrows. Genes in common in both enrichment analyses in Figure S3A and Figure S4 include upregulation of the tumor suppressors *Tmprss11e*, *Adamts18* and *Tgm3*, as well as induction of the pyroptosis regulating gene *GSDMc* and the epithelial-specific *Ets* transcription factor 1 (*Elf3*). Commonly down-regulated genes include activators of the lectin pathway of the complement system (*Fcna*), myosin light chain kinase 4 (*MLK4*), and pathogen recognition receptors (*Cd209*). These changes are plausible contributors to difference in tumor growth observed when Cxcr2 is targeted in melanocytes during transformation.

### CXCR2 Contributes to an Immunosuppressive Melanoma Tumor Microenvironment

While GSEA identified enrichment of stem cell and growth-associated gene sets in *Braf/Pten/Cxcr2^-/-^* mice, it also revealed induction of gene sets associated with CD4+ T cell activation, lymphocyte activation, and leukocyte proliferation (Figure 2H). These results prompted analysis of the immune cell infiltrate between *Braf/Pten/Cxcr2^WT^* and *Braf/Pten/Cxcr2^-/-^* tumor-bearing mice. We first utilized the murine Microenvironment Cell Population counter (mMCPcounter, (30)), an immune deconvolution algorithm developed for bulk murine RNA sequencing data. mMCPcounter predicted an increase in CD3+ T cells, CD8+ T cells, monocytes, lymphatic vessels, and eosinophils, as well as a decrease in mast cells, NK cells, and endothelial cells (p<0.05) (Figures 3A, S5A), suggesting enhanced anti-tumor immunity in the *Braf/Pten/Cxcr2^-/-^* TME. To analyze the immune environment *in vivo*, we defined the profile of CD45+ cells from *Braf/Pten/Cxcr2^WT^* and *Braf/Pten/Cxcr2*^-/-^ tumor-bearing mice using FACS analysis. In agreement with the mMCPcounter predicted leukocytic infiltrates, we observed that deletion of *Cxcr2* in melanocytes undergoing transformation skewed the TME toward anti-tumor immunity. FACS analysis of the CD45+ cells in the tumors of *Braf/Pten/Cxcr2*^-/-^ mice revealed a decrease in the immunosuppressive Ly6G+CD11b+ (p<0.01) and CD14+ G-MDSC (p<0.05) cells with no change in total CD11b+ cells (Figure 3B), in addition to a trend toward decreased CD25^hi^CD45+CD3+ regulatory T cells and a trend toward an increase in the frequency of CD3+CD8+T cells. There was also a significant increase in memory CD44+CD4+ T cells (p<0.05) and activated CD69+CD8+ T cells (p<0.05) within the *Braf/Pten/Cxcr2*^-/-^ tumors (Figure 3C, S5D). FACS analysis of peripheral blood cells revealed no significant change in any immune population (Figure S5B).

**Fig 3.**
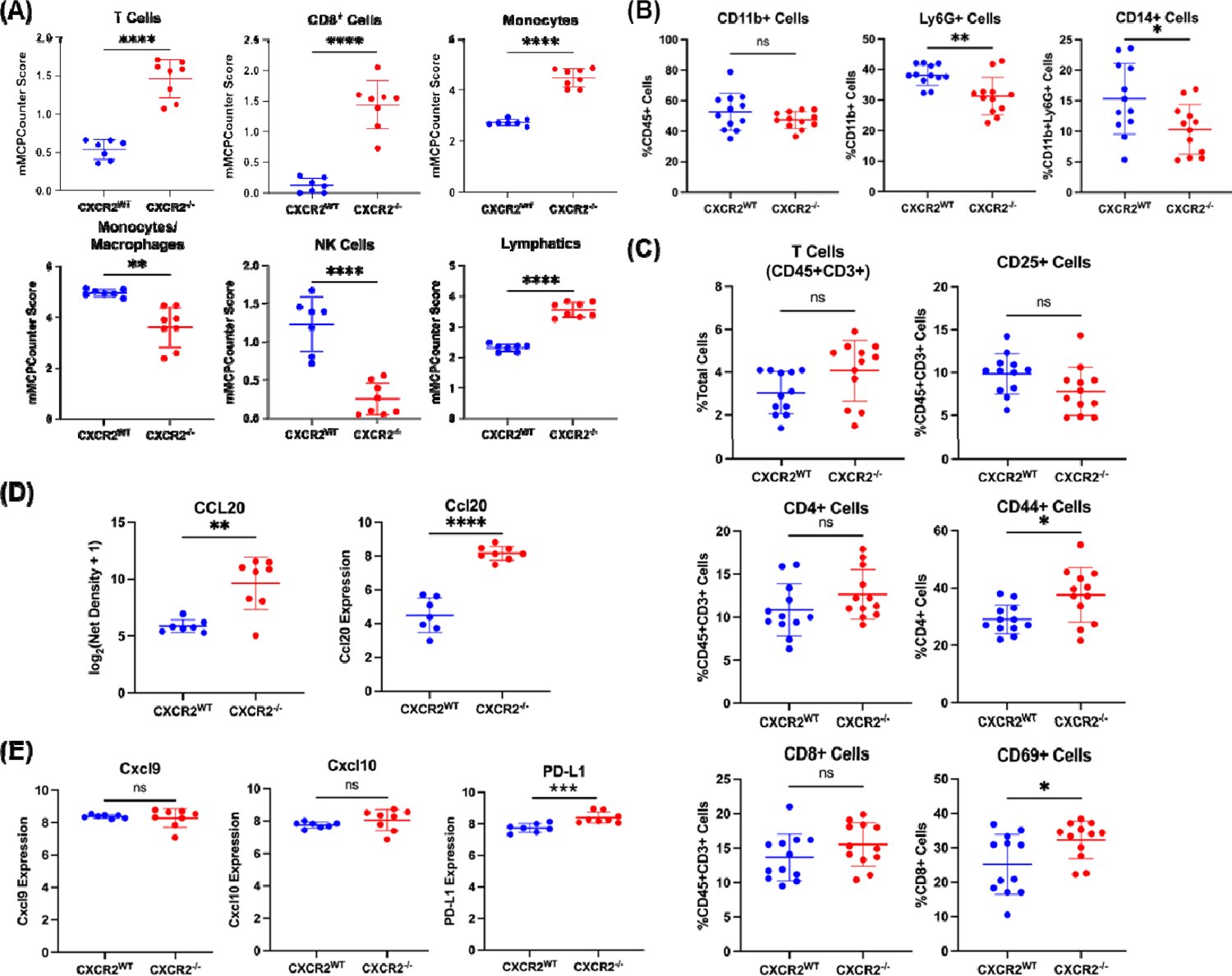
The immune infiltrate of *Braf^V600E^/Pten^-/-^* tumors is altered with loss of *Cxcr2*. **a** mMCPCounter analysis performed on bulk RNAseq data from *Braf^V600E^/Pten^-/-^*melanoma tumors with or without Cxcr2 predicts significantly enhanced infiltration of T cells, CD8+ T cells, monocytes, NK cells, and lymphatic vessels into *CXCR2^-/-^* tumors. **b** FACS analysis of CD45+ myeloid cells in *Braf^V600E^/Pten^-/-^* melanoma reveals decreased MDSC-like cells in *CXCR2^-/-^* tumors. **c** FACS analysis of CD45+ cells in *Braf^V600E^/Pten^-/-^* melanoma tumors identified changes in activated CD4+CD44+ T cells and CD8+CD69+ T cells. **d** Cytokine array for 62 cytokines expressed in TME of *Braf^V600E^/Pten^-/-^* tumors revealed one major cytokine, CCL20, that is strongly upregulated with loss of *Cxcr2* (n=4/genotype) based on net density. These data are complemented by increased *Ccl20* mRNA with loss of *Cxcr2* in *Braf^V600E^/Pten^-/-^* tumors. **e** *Cxcl9, Cxcl10*, and *PD-L1* expression based upon RNAseq analysis from *Braf^V600E^/Pten^-/-^* tumors expressing or not expressing Cxcr2 in melanocytes. All statistical significance determined via Welch’s t-test.

The identified differences in immune cell infiltrate are highly suggestive of altered cytokine signaling within the TME. Therefore, a 62-cytokine array was performed on *Braf/Pten/Cxcr2^WT^* (n=4) and *Braf/Pten/Cxcr2^-/-^* (n=4) tumor lysates. CCL20, an inflammatory chemokine that is highly chemotactic for CCR6-expressing lymphocytes and dendritic cells, is strongly upregulated (24-fold) in the *Braf/Pten/Cxcr2^-/-^* TME (Figure 3D). In addition, RNAseq analysis revealed a significant increase in PD-L1 expression in tumors from *Braf/Pten/Cxcr2^-/-^* mice compared to *Braf/Pten/Cxcr2^WT^* mice (Figure 3E). Furthermore, M-CSF, eotaxin, and MIP-2 were slightly increased, which could contribute to myeloid cell infiltration, and there was a slight decrease in IL-1β in the tumors from *Braf/Pten/Cxcr2^-/-^* mice as compared to tumors from *Braf/Pten/Cxcr2^WT^* mice (Figure S5C). These data suggest that targeted deletion of *Cxcr2* in melanocytes during tumorigenesis results in a marked increase in *Ccl20* and additional subtle changes in the cytokine milieu of the TME.

### CXCR1/CXCR2 Antagonist SX-682 Inhibits *Braf^V600E^/Pten^-/-^* and *NRas^Q61R^/Ink4a^-/-^* Melanoma Tumor Growth and Promotes Anti-Tumor Immunity

Having established the importance of *Cxcr2* in the development, growth, and TME of *Braf/Pten* melanoma tumors, we sought to evaluate the therapeutic potential of systemic Cxcr1/Cxcr2 inhibition. Thus, chow containing the Cxcr1/Cxcr2 antagonist SX-682 (31) was administered to four-week-old mice. After two weeks of eating vehicle control or SX-682 containing chow, 4-HT was applied to the backs of the mice for three successive days. Following a month of continuous feeding on control or SX-682-containing chow, we observed that mice fed SX-682-containing chow exhibited a trend toward reduction in tumor volume compared to mice fed vehicle control chow (Figure 4A, p=0.07, 802.5±724.01mm^3^ for control; 230.20±373.21 mm^3^ for SX-682). Moreover, there was a trend toward decreased tumor formation in SX-682-fed mice (p=0.145), where only 40% of SX-682-fed mice developed tumors compared to 75% of control-fed mice (Figure 4B). Similarly, *NRas^Q61R^/Ink4a^-/-^* (*NRas/Ink4a)* mice were fed chow containing SX-682 or control chow, and tumors that developed over five months were counted and measured. We observed that SX-682 treatment significantly suppressed tumor growth (p=0.041, Figure 4C) but only trended toward a decrease in tumor incidence (p=0.111, Figure 4D).

**Fig 4.**
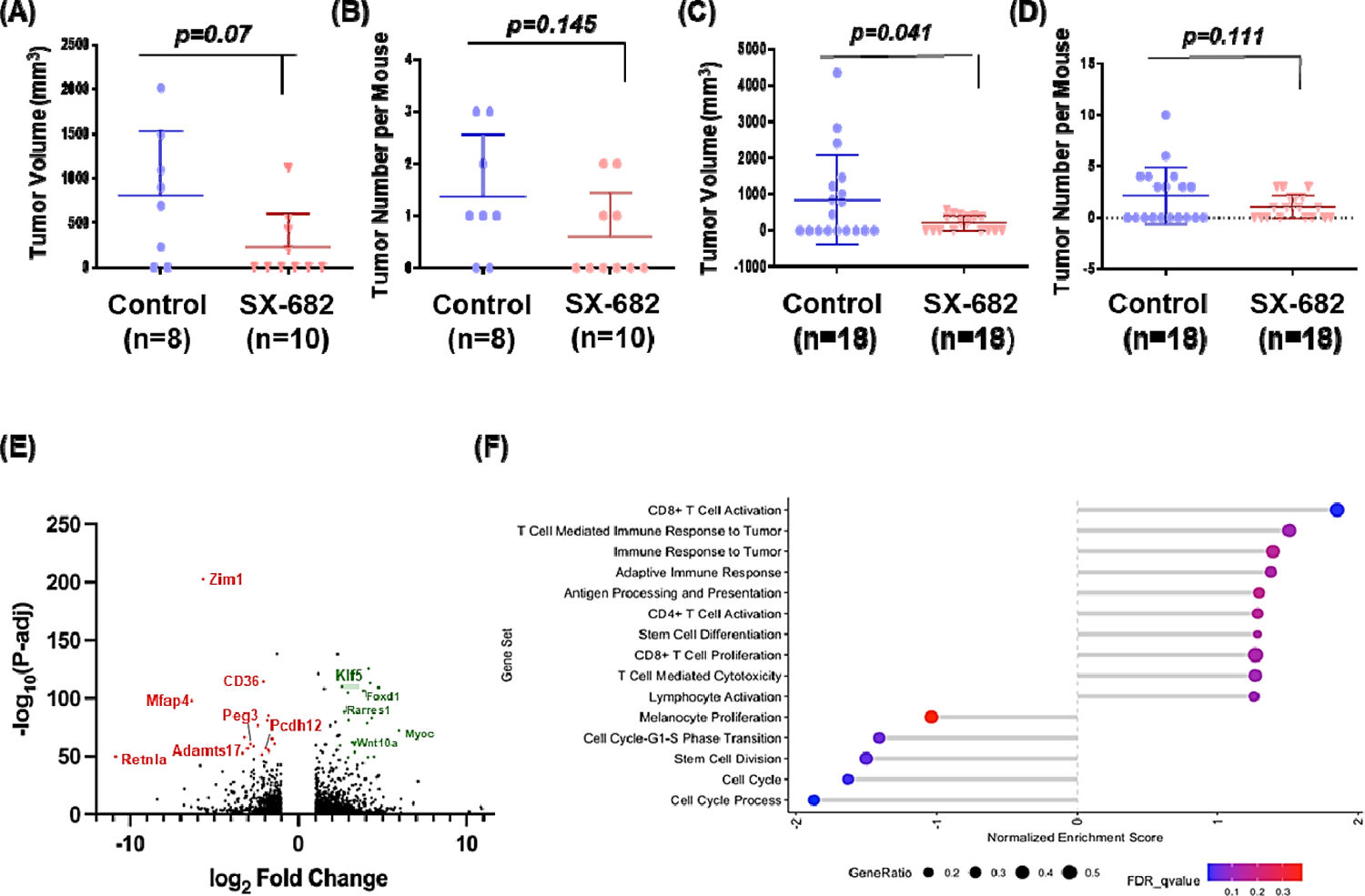
SX-682 affects *Braf^V600E^/Pten^-/-^* and *NRas^Q61R^/Ink4a^-/-^* tumorigenesis. **A,b** *Braf^V600E^/Pten^-/-^* and **c,d** *NRas^Q61R^/Ink4a^-/-^* mice were fed chow containing SX-682 or vehicle continuously through tumor formation, and tumors were measured and counted. Significance was determined using a Welch’s t-test. **e** A volcano plot showing fold change and significance of differential gene expression between tumors from SX-682-fed and control-fed *Braf^V600E^/Pten^-/-^* mice. **f** Gene set enrichment analysis of SX-682 treated or control *Braf^V600E^/Pten^-/-^* tumors identifies gene sets enriched in SX-682 treated tumors (positive normalized enrichment score) or enriched in control tumors (negative normalized enrichment score).

RNA sequencing analysis of control and SX-682 treated tumors from *Braf/Pten* mice identified nearly 3000 differentially expressed genes with many trends similar to those observed in *Braf/Pten/Cxcr2^-/-^* tumors. A volcano plot shows that a significant number of genes were strongly up or down-regulated (log_2_ fold change of > 3) with a very high level of significance (-log_10_(P-adj)>50) (Figure 4E). Upregulated genes include those involved in regulation of growth, proliferation, and cell cycle; tumor suppression; differentiation/stemness; immune regulation; and motility and adhesion. Genes downregulated in response to Cxcr1/Cxcr2 antagonism with SX-682 include those involved in cell adhesion and cell proliferation, cell cycle and growth (Figure S3B).

GSEA of the tumors from *Braf/Pten* mice treated with SX-682 revealed significant increases in CD8+ T cell activation, with trends toward increased T cell-mediated immune response to the tumor, immune response to tumor, adaptive immune response, antigen processing and presentation, CD4+T cell activation, stem cell differentiation, CD8+ T cell proliferation, T cell-mediated cytotoxicity, and lymphocyte activation. There were significant decreases in genes involved in melanocyte proliferation, cell cycle process, cell cycle, stem cell division, and cell cycle G1-S transition (Figure 4F). mMCPcounter analysis of the tumor RNAseq data predicted an increase in CD8+ T cells (Figure 5A) and monocytes (Figure S6A), and a decrease in B-derived cells and cells of the lymphatics (p<0.01) in tumors from the SX-682-treated *Braf/Pten* mice (Figure S6A).

**Fig 5.**
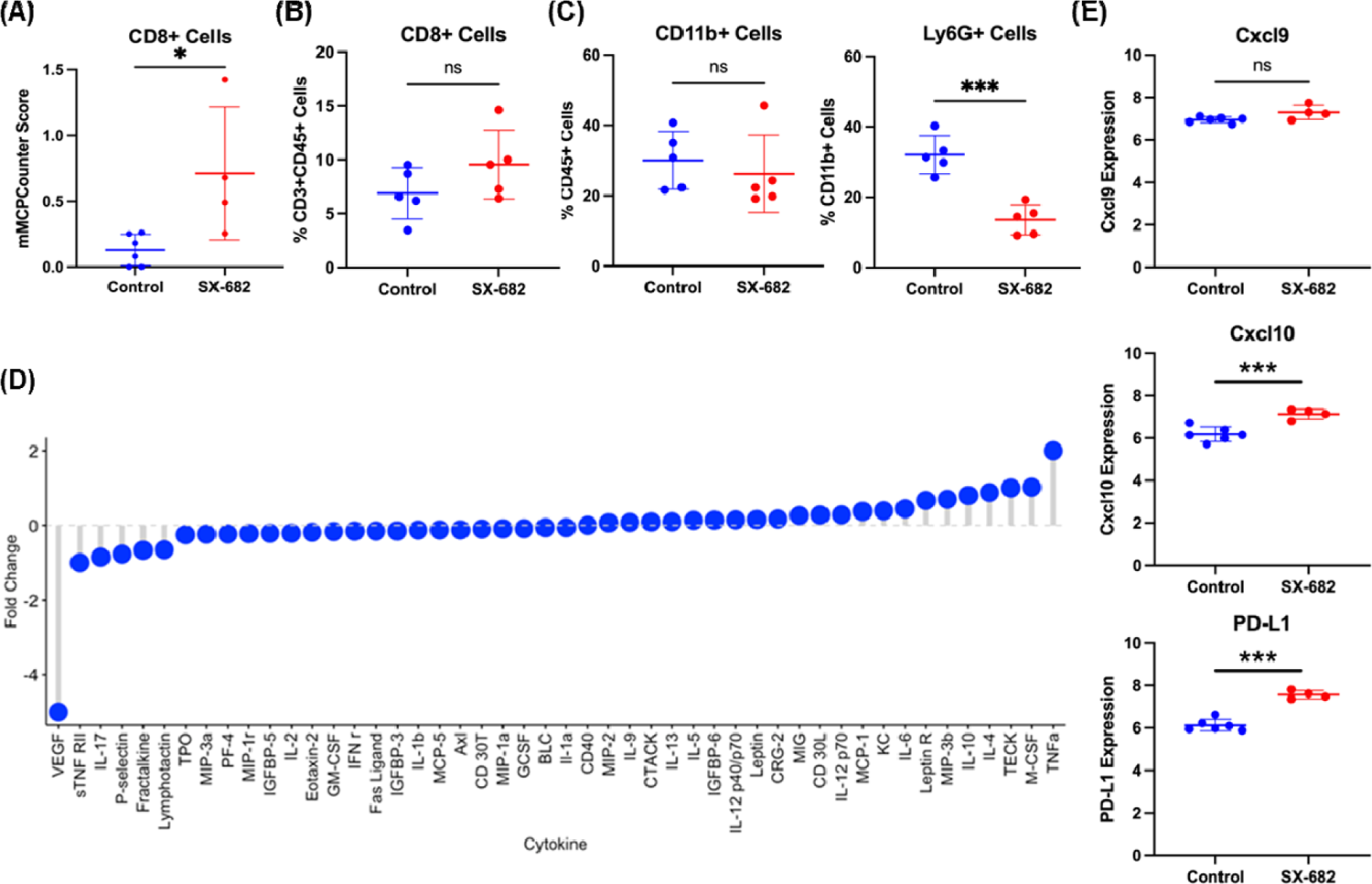
SX-682 alters the immune profile of *Braf^V600E^/Pten^-/-^* melanoma. **a** mMCPCounter analysis of bulk RNAseq data predicts enrichment for CD8+ T cell infiltrate into tumors following treatment with SX-682 (p<0.05). **b** FACS analysis confirms a trend toward increased CD8+ T cells in SX-682 treated *Braf^V600E^/Pten^-/-^* melanoma. **c** FACS analysis of CD45+ myeloid cells indicated a significant decrease in immunosuppressive CD11b+Ly6G+ cells, but no change in total CD11b+ cells. **d** A cytokine array was performed on control and SX-682 treated tumors, identifying a notable decrease in Vegf and an increase in Tnfα. **e** *Cxcl9, Cxcl10,* and *Pd-l1* expression based upon RNAseq analysis from SX-682 or control treated tumors. All statistical significance determined via Welch’s t-test.

FACS analysis of SX-682 treated *Braf/Pten* tumors revealed a trend toward increased CD8+ T cells (p=0.17), no change in CD11b+ cells, and a significant decrease in CD11b+Ly6G+ cells (p<0.001) (Figures 5B, C). Additional FACS analysis of tumor CD45+ cells showed a decrease in CD4+CD3+ cells (p<0.05) in tumors from the SX-682 chow-fed mice (Figure S6C). In peripheral blood, there was a significant decrease in CD44+ CD4+ T cells and CD62L+ CD4+ T cells and a trend toward increased CD69+ CD8+ T cells from mice fed SX-682 chow (p=0.059; Figure S6B). In addition, a cytokine array of tumor lysates (n=4 for each genotype) revealed a marked reduction in *Vegf*, indicating a reduction in tumor angiogenesis, and an increase in *Tnf*α, indicating a more inflammatory tumor microenvironment (Figure 5D). Moreover, RNAseq analysis of *Braf/Pten* tumors revealed that SX-682 induces expression of *Cxcl9, Cxcl10*, and *Pd-l1* (Figure 5E). Altogether, these data indicate that SX-682 alters the TME to stimulate anti-tumor immunity and reduce tumor growth.

### SX-682 Treatment of Melan-A, B16F0, and B16F10 Cells Reveals Tumor Cell-Specific Gene Modulation

Our murine experiments involved bulk RNA sequencing of tumors that contain tumor cells in addition to stromal and immune cells. To identify the specific effect of SX-682 treatment on tumors without the contribution of other cell types, we investigated the effect of SX-682 on non-tumorigenic Melan-A cells, tumorigenic B16F0 cells, and metastatic B16F10 cells *in vitro*. First, we evaluated Cxcr2 expression and found that B16F0 and B16F10 cells express significantly more Cxcr2 than Melan-A cells, as evaluated by mRNA levels and surface protein labeling (Figure 6A, B). We then μM) on the growth of these cells and observed that SX-682 treatment resulted in a small but significant inhibition of growth in B16F0 and B16F10 cells *in vitro* based on the percentage of cells staining positively for KI-67 (Figure 6C) and cell number (Figure S7A). In addition, SX-682 treatment of B16F0 and B16F10 cells *in vitro* also reduced production of both Cxcl1 (KC) and vascular endothelial growth factor (Vegf) as evaluated by cytokine array (Figure 6D), again indicating the potential for SX-682 to impact the immune profile of the TME.

**Fig 6.**
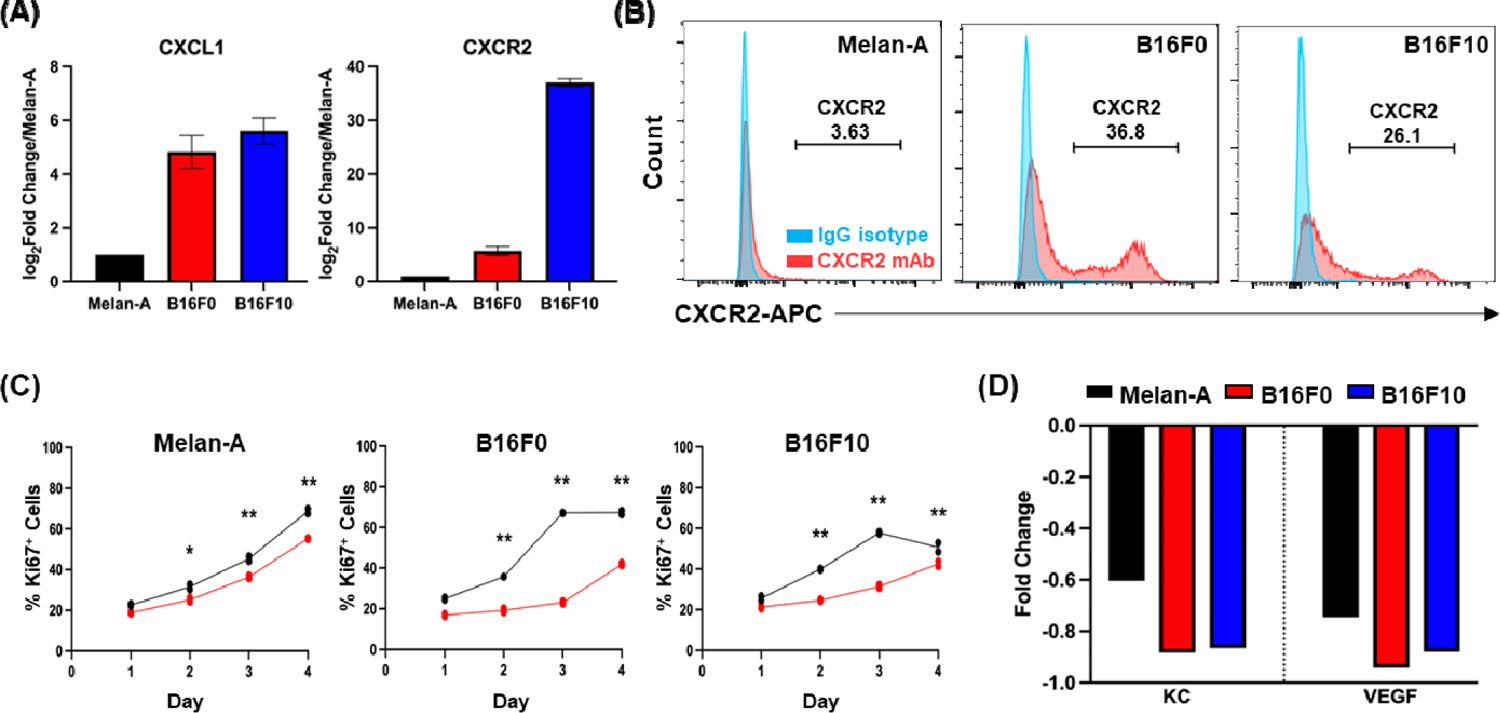
Tumor cell-specific impacts of SX-682. CXCL1 and CXCR2 expression on Melan-A, B16F0, and B16F10 cells based on **a** the NCBI database and **b** CXCR2 expression in Melan-A, B16F0 and B16F10 cells based on flow cytometry. **c** Cell lines were treated with 5 µM SX-682 (or DMSO control) for 4 days prior to staining with Pacific Blue-Ki67 for FACS analysis. The percentage of positive staining cells was significantly decreased in the SX-682 treated cells for all cell lines (analyzed using a two-way ANOVA with Benjamini and Hochberg (BH) correction for multiple tests). **d** Cytokine array of SX-682-treated Melan-A, B16F0 and B16F10 cells shows that SX-682 treatment reduced the expression of KC and VEGF in all three cell lines.

To identify tumor cell-specific transcriptional changes following SX-682 treatment, we performed RNA sequencing on each of the three cell lines. Of the total differentially expressed genes, expression of 4024 genes was altered in all three lines. An additional 860 genes were differentially expressed in both tumorigenic B16F0 and B16F10 lines in response to SX-682 (Figure S3C and S7B). Commonly upregulated genes include those involved in apoptosis and cell stress response and suppression of gluconeogenesis. In contrast, commonly down-regulated genes include those involved in methylation, RNA splicing, and cell cycle processes (Figure S3C and S7B). Reverse phosphoprotein analysis (RPPA) identified SX-682-induced decreases in phosphoproteins involved in growth (Akt, Braf, pS445-Braf, Cdc2-pY15, Cdc6, Gsk-3b, mTor, mTor pS2448, Mmp14, Pax8, and S6), as well as SX-682-induced increases in immunomodulatory proteins (Sting, Pd-1, Pd-l1, Trim25, and Annexin I); proteins involved in the regulation of apoptosis (Puma, Blc2, Bcl2A1, BclxL, Smac); tumor suppressors (Tsc2, Wtap); and cell cycle regulators (Cdc25, Cdc42, Plk1, Egfr, β-catenin expression is increased following SX-682 treatment. This is counter-intuitive for SX-682 inhibition of tumor growth, as the Wnt/ -βcatenin pathway often drives melanoma tumor growth and metastasis. However, we observed that the phosphorylated forms of β ubiquitin-mediated degradation are increased as well. This indicates that β marked for degradation, thus diminishing the potential for enhanced tumor growth. There were also increases in proteins involved in motility: myosin-Iia, Pak, Cdc-42, myosin Iia_pS1943, and Hmha1 (Figure S7C, D). Finally, there were only subtle changes in cytokine expression in response to SX-682 treatment *in vitro,* and these were inconsistent across the three cell lines (Figure S7E). Altogether, these results suggest that multiple compounding signals are induced in cells treated with SX-682, including a decrease in growth signaling, modulation of apoptosis, enhanced anti-tumor immunity, and altered cell cycle processes.

### *Tfcp2l1* distinguishes the Cxcr2^WT^ from the Cxcr2 Perturbed Phenotype

To better understand the complex transcriptional reprogramming that occurs when CXCR2 activity is diminished via knockout or with SX-682 treatment, we compared differentially expressed genes in *Braf/Pten/Cxcr2^-/-^* tumors, SX-682-treated tumors, and SX-682-treated tumorigenic B16F0 and B16F10 cell lines compared to controls. We noted that based upon our search for genes with a minimum of a log2 fold change >2 and a p-value <0.05, only one gene stood out as significantly upregulated across all four models compared to the respective controls: Transcription factor CP2 like-1(*Tfcp2l1)* (Figure 7A, B). To verify the RNA sequencing results, we performed RT-PCR analysis of RNA samples from MelanA, B16F0, and B16F10 cells to determine *Tfcp2l1* expression. With this assay, we show that SX-682-treatment elevates *Tfcp2l1* expression in the tumorigenic cell lines (Figure S9).

**Fig 7.**
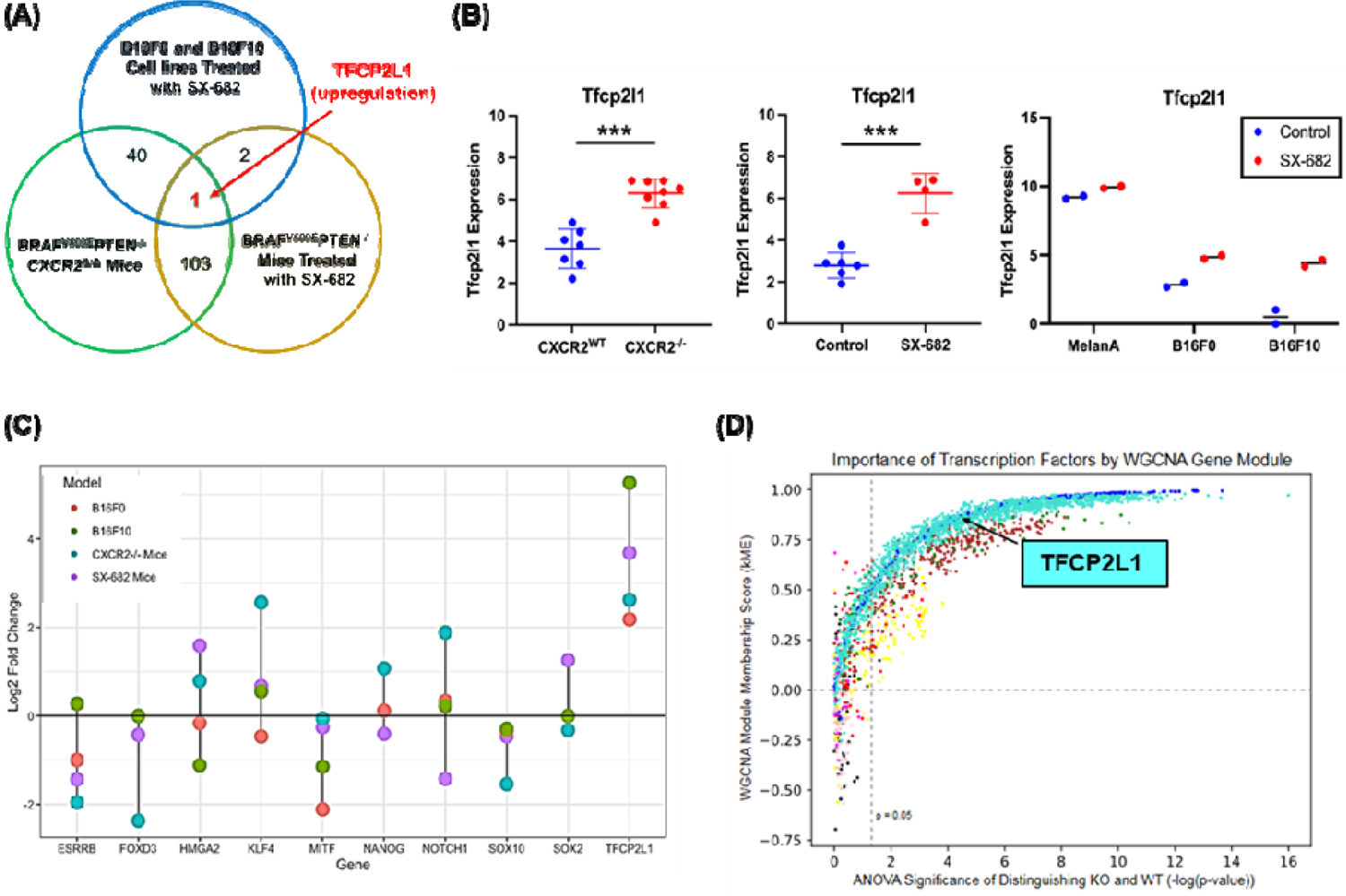
*Tfcp2l1* is commonly upregulated across three models of CXCR2 perturbation. **a, b** In comparing expression data from *Braf^V600E^/Pten^-/-^/Cxcr2^-/-^* tumors. *Braf^V600E^/Pten^-/-^* tumors treated with SX-682, and B16F0 and B16F10 cell lines treated with SX-682, *Tfcp2l1* was consistently upregulated compared to appropriate controls (as determined by Welch’s t-test). **c** Log2 change for *Tfcp2l1* and related genes across experimental groups based upon RNAseq anal fold sis. **d** Identification of transcription factors central to Weighted Correlation Network Analysis (WGCNA) co-expressed gene modules (by kME) and significantly differentially expressed between *Braf^V600E^/Pten^-/-^/Cxcr2^-/-^* and *Braf^V600E^/Pten^-/-^/Cxcr2^WT^* tumors. TFs are colored by gene module and show varying levels of centrality to each module and importance in distinguishing WT and KO tumors. Turquoise dots represent transcription factors that are up in the *Braf^V600E^/Pten^-/-^/Cx*r2*^-/-^* tumors and blue dots represent transcription factors that are up in the *Braf^V600E^/Pten^-/-^/Cxcr2^WT^* tumors.

Tfcp2l1 is a transcription factor that contributes to the maintenance of stemness in pluripotent stem cells and can also exhibit tumor suppressive activity (32, 33, 34, 35, 36). The Krupple-like Factor (KLF) family of transcription factors works with and can be induced by Tfcp2l1 to modulate induction and maintenance of naïve pluripotency in mouse primordial germ cells (37, 38, 39). It has been previously reported that *Tfcp2l1* is positively associated with expression of pluripotency genes including *Nanog, Oct4, Sox2,* and *Esrrb* in mouse embryonic stem cells (42). However, our data suggest a complex relationship between Cxcr2 perturbation and Tfcp2l1-related gene expression. In the *Braf/Pten* model, stemness marker *Esrrb* and neural crest markers *Foxd3* and *Sox10* were decreased when Cxcr2 was deleted. In contrast, stemness markers *Tfcp2l1*, *Klf4* and *Hmga2,* were increased. In SX-682 treated *Braf/Pten* model, there was a trend toward a decrease in stemness marker *Esrrb,* a significant decrease in the neural crest marker *Sox10*, and a small but significant decrease in the melanoblast marker *Mitf* (Figures 7C, S8A-L). The melanocyte differentiation marker Tyr was increased in both the CXCR2^-/-^ and the SX-682 treated *Braf/Pten* mouse models. In the B16F0 and B16F10 cells, RT-PCR analysis revealed that stemness markers *Esrrb, Hmga2, Myc, Sox2*, neural crest marker *Sox10,* and melanoblast marker *Mitf* were significantly decreased in response to SX-682-treatment *in vitro*. *Foxd3* was significantly decreased in B16F10 and trended toward significant reduction in B16F10. In contrast, stemness markers *Tfcp2l1, Nanog* and *Notch1* were increased, while there was a decrease in *Tyr* expression (Figure S9). Altogether, these data imply that with ablation of Cxcr2 activity, there is an increase in some stemness markers, a decrease in neural crest markers, and a trend toward a decrease in melanoblast markers. However, there is variability in the mix of these markers from model to model. In the mouse models, tyrosinase (*Tyr*) continues to be highly expressed, though in the B16 cell cultures, SX-682 decreased its expression.

To support the relevance of upregulation of the transcription factor, *Tcfp2l1*, in association with loss of CXCR2 signaling, we performed an orthogonal approach based on weighted gene co-expression network analysis (WGCNA). WGCNA was applied to the RNA-seq data from these tumors to generate groups of highly correlated genes, or gene modules, that functionally distinguish *Braf/Pten/Cxcr2^-/-^* and *Braf/Pten/Cxcr2^WT^* tumors (Figure S10). Using an ANOVA test between sample conditions, we found six distinct modules that significantly distinguish the transcriptional programs of *Cxcr2^WT^* and *Cxcr2^-/-^* tumors (Figure S10). Gene ontology (GO) analysis showed each module is enriched in distinct functions: the *Cxcr2^WT^*-upregulated modules are enriched in GO terms such as protein localization to mitochondrion (blue), aerobic respiration and oxidative phosphorylation (green and brown, respectively), and signaling (yellow), while the *Cxcr2^-/-^*-upregulated modules are enriched for GPCR signaling (red) and skin development (turquoise,). These changes in gene expression may result as an adaptation to the loss of CXCR2 function. Interestingly, the WGCNA module membership score (kME) indicated that Tfcp2l1 is central to the turquoise module (kME = 0.854) and significantly upregulated in the *Cxcr2^-/-^* samples (FDR-adjusted p=0.0000286) (Figure 7D).

Finally, to define the activity of Tfcp2l1 following CXCR2 perturbation, we performed chromatin immunoprecipitation and sequencing analysis (CHIPseq) on B16F0 tumorigenic melanoma cells following treatment with vehicle or SX-682. In identifying promoters bound by Tfcp2l1 in each condition in addition to RNAseq data, we can delineate SX-682-induced gene set enrichment. Interestingly, enrichment analysis of Tfcp2l1-bound genes revealed that SX-682 treatment increased expression of genes associated with the adaptive immune system and response to hormones (Figure S11A). SX-682 treatment also enriched Tfcp2l1 binding to and repression of genes involved in β-catenin independent Wnt signaling pathways, and catabolism (Figure S11B). When data from RNAseq, RPPA, and ChIPseq analysis were examined using Metascape, key regulatory pathways emerged as commonly associated with CXCR2 loss of function (Figure S11C, D). These data are consistent with the observed reduction in tumor growth when CXCR2 signaling is blocked and suggest that changes in gene expression are associated with Tfcp2l1 transcriptional control.

### Does CXCL1 activation of normal melanocytes suppress the TFCP2L1 transcriptional program?

To gain insight into how CXCL1 activation of CXCR2 regulates the expression of stemness and differentiation markers, RNAseq analysis was performed on normal human epidermal melanocyte (NHEM) cultures treated with CXCL1 or with CXCL1 and SX-682 (Figure S12). CXCL1 supplementation was utilized in this model to recreate the enhanced baseline CXCR2 activation of tumorigenic cells. Consistent with this, CXCL1 increased the proliferation of NHEM cells *in vitro* (Figure S12A). Moreover, CXCL1 treatment increased expression of a number of genes, and this effect was lost with SX-682 treatment (Figure S12B, white oval). For example, CXCL1 treatment of NHEM cells induced a trend toward increased expression of *MITF, BMP6, WNT5A, and SOX10*, and the addition of SX-682 reversed this trend. Moreover SX-682 treatment resulted in induction of expression of a host of genes that are lowly expressed in control and CXCL1 treated NHEM cells (Figure S12B, yellow box) and suppresses expression of many highly expressed genes in control and SX-682-treated NHEMs (Figure S12B, orange box). SX-682 also induced a trend toward elevated *TFCP2L1, KLF4, FOXD3, FOXD1*, and *CCL20* expression over that produced by CXCL1 alone (Figure S12C). These data clearly show that loss of CXCR2 activity dramatically alters gene expression, resulting in reduced CXCL1-induced proliferation of NHEM. In addition to the effects on stemness and differentiation markers, we also found that several chemokines, interleukins (Table S1), and TNF-related cytokines and interferons (Table S2) were altered when NHEMs were treated with combined CXCL1 and SX-682. SX-682 treatment increased expression of inflammatory genes *CCL20, IL18R1, IL1RL1* and decreased expression of chemokines associated with macrophage and MDSC recruitment (*CCL2, CCL7, CCL8, CXCL1, CXCL12, CXCL6* and *IL33)* as well as TNF family members involved in MAPK activation, osteoclastogenesis, and B cell activation (*C1QTNF2, TNFRSF21, TNFSF11*, and *TNFSF13B)*.

Taken together, our data from both tumor models and *in vitro* studies show that CXCR2 activation is associated with activation of the MAPK cascade, AKT, and WNT signaling, expression of chemokines that recruit MDSCs and protumor macrophages, and enhanced tumor growth. In contrast, loss of CXCR2 or inhibition of CXCR1/CXCR2 in melanoma progenitor cells is associated with expression of genes associated with inflammation, T cell recruitment, pluripotency, and reduced tumorigenicity. The mechanism for these changes in gene expression are in part due to induction of *Tfcp2l1*, a transcription factor that regulates genes that suppress tumorigenicity.

### Discussion

The CXCR1 and CXCR2 receptors are G protein-coupled receptors that generate downstream signals including PI3K and AKT, often implicated in growth (6, 11, 40, 41, 42). The role of CXCR2 in cell motility has been well characterized, and the signals generated through this receptor leading to activation of AKT and ERK also modulate cell proliferation and growth(43, 44).

CXCR1 has been reported to be important for the renewal of a population of stem cell-like cells in human breast cancer (45). In mice, CXCR2 controls functions normally regulated by CXCR1 in humans, thus it is plausible that CXCR2 may also modulate stemness. Here, we examined the role of CXCR2 in melanocyte tumorigenesis and observed that loss of CXCR2 in tyrosinase-expressing melanocytes reduced melanoma tumor burden in *Braf/Pten and NRas/Ink4a* murine melanoma and modulated the expression of melanocyte stemness and differentiation markers.

We observed that the mechanism by which loss of Cxcr2 activity during melanocyte tumorigenesis resulted in reduced tumor growth in *Braf/Pten* mice was due to major changes in gene expression, with decreased expression of genes involved in proliferation and increased expression of genes associated with tumor suppression, T cell recruitment and differentiation, and apoptosis. These gene expression data from RNAseq analysis were further supported by phospho-proteomic data. We observed that loss of Cxcr2 activity in tumor cells resulted in a change in the tumor immune microenvironment, with increased CD8+ T cells and reduced macrophages and MDSC-like cells. When Cxcr1/Cxcr2 were antagonized in *Braf/Pten* mice and tumorigenic melanoma cell lines via treatment with SX-682, similar alterations in the gene expression profiles were achieved, and this was accompanied by development of anti-tumor immune microenvironment.

When we looked for genes significantly induced in Cxcr2^-/-^ tumors, SX-682 treated tumors, and B16F0 and B16F10 cell lines, one common gene emerged: *Tfcp2l1*. *Tfcp2l1* is a crucial transcription factor that induces the expression of genes associated with stemness in embryonic stem cells (32). As such, we probed the relationship between Tfcp2l1, differentiation along the melanocyte lineage, and cancer stem cells within melanoma.

Much of our understanding of melanocyte lineage came from *in vitro* studies that involved the differentiation of human pluripotent stem cells along a neural crest lineage, then on to form melanocytes (46). Wnt ligands and Bmp4 induce the early transition of Oct4+Nanog+ pluripotential cells into Sox10+ neural crest cells. Exposure to endothelins and Bmp4 promotes neural crest cell differentiation to Mitf+cKit+ melanoblasts, and these can be terminally differentiated to Tyr+Oca2+ melanocytes through continued exposure to Wnt ligands, Bmp4, and induction of intracellular cAMP (47). In the melanoma models used in our studies, the targeted alterations in gene expression (*Braf/Pten/Cxcr2^-/-^*) occur in tyrosinase expressing melanocytes. Interestingly, while loss of CXCR2 expression or activity was not associated with reduction in tyrosinase in our mouse models, we noted a decrease in the expression neural crest markers *Sox10* and *Foxd3* in tumors that developed when Cxcr2 activity was ablated. In addition, there was an increase in expression of some markers associated with pluripotency or stemness.

While we do see consistent *Tfcp2l1* induction across all our models of Cxcr2 perturbation, trends in Tfcp2l1-regulated genes are not as clear. There is a trend toward increased *Klf4, Hmga2, Notch1, Myc*, and *Stat3* expression which would suggest that tumors with loss of Cxcr2 are less differentiated. However, Esrrb, which has been established as a direct target of Tfcp2l1 binding and induction in ESCs (39), is significantly decreased in our *Cxcr2^-/-^* tumors. The implications of this shift in stemness markers in relation to melanoma aggression, treatment sensitivity, and overall prognosis is currently unknown.

Our finding that loss or inhibition of Cxcr2 activity in melanocytic cells results in changes in markers associated with stemness, neural crest cells, and melanoblasts in association with a reduction of tumor formation and growth is somewhat paradoxical. However, human melanoma tumors are quite heterogeneous (48), with stem-like cell populations as well as more differentiated populations expressing MITF, TYR, and MELANA. Of note, nests of stem-like melanoma cells have been identified in metastatic lesions in head and neck cancer patients and shown to express *NANOG, OCT4, SOX2, KLF4,* AND *cMYC* (48). Moreover, melanocytes and melanoma cells have been dedifferentiated to iPSCs by transfecting in *Oct4, c-Myc,* and *Klf4* expression vectors. The resulting iPSCs express *Nanog* and *Oct4* and can be differentiated into fibroblast-like cells (49). Our data suggest that loss of CXCR2 signaling may reduce sub-populations of melanoma cells expressing the neural crest marker *Sox10* and stem cell marker *Esrrb* but increases populations with the stemness markers *Klf4, Hmga1,* and *Tfcp2l1*. Moreover, the gene expression pattern in the six functionally enriched states of tumor cells previously established by single-cell transcriptomics: melanocytic, neural crest-like, antigen-presenting, RNA processing, stem-like, and stress-like appear to be altered with loss of Cxcr2 signaling, especially in the melanocytic state (50).

## Conclusion

We demonstrate that targeted deletion of *Cxcr2* in tyrosinase-expressing melanoma precursor cells concurrent with induction of the *Braf^V600E^* transgene and loss of *Pten* expression or induction of *NRas^Q61R^* and loss of *Ink4a*, resulted in a significant reduction of melanoma burden. Notably, we also observed reduced expression of genes involved in growth, increased expression of genes involved in tumor suppression, and promotion of an anti-tumor immune environment when *Cxcr2* was deleted in tyrosinase-expressing melanoma precursor cells during transformation. Importantly, we show that the CXCR1/CXCR2 antagonist, SX-682, accomplishes a similar reduction in melanoma tumor burden, establishes an anti-tumor immune microenvironment, and significantly alters the transcriptional profile of melanoma cells when delivered during the transformation process. A key mechanism for these transcriptional changes involves increased expression of *Tfcp2l1*, a predicted tumor suppressive transcription factor when Cscr2 activity is blocked.

Our data support combining CXCR1/CXCR2 antagonists with immunotherapy for melanoma patients. Consistent with this concept, we have shown that the antagonism of Cxcr2 upregulates PD-L1 expression and enhances the response of melanoma cells to anti-PD-1(9). Moreover, CXCR1/CXCR2 antagonists combined with anti-PD-1 are currently in clinical trials for the treatment of melanoma (NCT03161431). Moving forward, it will be essential to identify the subset of patients most likely to respond to this combination therapy and to develop protocols for maximal response.

## Supporting information

All Supplemental Material

## List of abbreviations

CXCR2: C-X-C Motif Chemokine Receptor 2.

Ink4a: inhibitor of cyclin-dependent kinase 4a.

Pten: phosphatase and tensin Homolog.

TFCP2L1: Transcription Factor CP2 Like 1.

MDSC: myeloid-derived suppressor cells.

TME: tumor microenvironment.

PI3K: phosphatidylinositol-3-kinase.

MAPK: mitogen-activated protein kinase.

AKT: protein kinase B.

NF-κB: nuclear factor kappa-light-chain-enhancer of activated B cells.

KC: keratinocyte chemoattractant.

MIP-2: macrophage-inflammatory protein-2.

LIX1: limb and CNS expressed 1.

TCGA: the cancer genome atlas.

GEPIA: gene expression profiling interactive analysis.

GSEA: gene set enrichment analysis.

WGCNA: weighted gene co-expression network analysis.

PD-1: programmed cell death protein 1.

CRE: cis regulatory element.

4-HT: Hydroxytamoxifen.

mT/mG: a cell membrane-localized Tomato (mT) and EGFP (mG) as a two-color fluorescent Cre-reporter allele.

RNAseq: RNA sequencing.

Tmprss11e: transmembrane serine protease 11e.

Adamts18: ADAM metallopeptidase with thrombospondin type 1 motif 18.

Tgm3: transglutaminase 3.

GSDMc: gasdermin C.

Elf3: E74 like ETS transcription factor 3.

Fcna: ficolin A.

MLK4: myosin light chain kinase 4.

FACS: fluorescence activated cell sorting.

M-CSF: macrophage colony-stimulating factor.

VEGF: Vascular endothelial growth factor.

RPPA: reverse phosphoprotein analysis.

mTOR: mammalian target of rapamycin.

MMP: matrix metalloproteinase.

PAX8: paired box gene 8.

STING: stimulator of interferon genes.

TRIM25: tripartite motif containing 25.

PUMA: p53 upregulated modulator of apoptosis.

BclxL: B-cell lymphoma-extra large.

Smac: second mitochondrial activator of caspases.

TSC2: tuberous sclerosis complex 2.

WTAP: Wilms tumor suppressor 1 associated protein.

PLK1: polo like kinase 1.

PRAS40: the proline-rich AKT substrate of 40 kDa.

HMHA1: minor histocompatibility protein HA-1.

KLF: krupple-like factor.

Nanog: nanog homeobox.

Oct4: octamer-binding transcription factor 4.

Sox2: SRY-box transcription factor 2.

Esrrb: estrogen related receptor beta.

Notch1: notch receptor 1.

Hmga2: high mobility group AT-hook 2.

Foxd3: forkhead box D3.

ANOVA: analysis of variance.

CHIPseq: chromatin immunoprecipitation and sequencing analysis.

NHEM: normal human epidermal melanocyte.

MELANA: melanocyte antigen.

## Declarations

### Ethical approval and consent to participate

Animal studies were approved by the Vanderbilt Institutional Care and Animal Use Committee (IACUC) and were performed in accordance with Vanderbilt IACUC guidelines.

### Consent for publication

Not applicable.

## Acknowledgments

We thank Dorothea Bennett for the MelanA cell line (University of Texas) and Christine Burd (The Ohio State University School of Medicine) for the *Tyr-CRE-ERT2-NRas^Q61^R/p16Ink4a*^-/-^ mice. We appreciate Tracy Handel (University of California, San Diego) for her helpful comments during the preparation of this manuscript.

## Author Contributions

J Yang performed the animal experiments, K Bergdorf analyzed the RNAseq data, C Yan analyzed human datasets from multiple sources, e.g., GEO, TCGA, TIDE, Riaz et al., 2017, and Chen et al., 2016, S-C Chen and D Ayers performed the biostatistical analysis, Q Liu, X Liu extracted the raw RNAseq data and provided analysis, W Luo helped with the ChIP-seq experiments, M Boothby provided immunology expertise, S M Groves, AN Oleskie, and V Quaranta assisted with the transcription factor analysis, JA Zebala and DY Maeda provided expertise for SX-682 experiments, A Richmond designed the study, oversaw the gathering and interpretation of data, all the authors contributed to the writing of the manuscript.

## Availability of data and materials

The datasets supporting the conclusions of this article are available in the Gene Expression Omnibus under accession GSE223290.

