## Supplementary material for "CXCR2 expression during melanoma tumorigenesis controls transcriptional programs that facilitate tumor growth": All Supplemental Material

**Supplementary Methods:**

**Analysis of patient sequencing data.**

RNA-Seq analysis of CXCL1, 2, 3, 5, and 8 from utilized patient nevi and melanoma samples available at NCBI Gene Expression Omnibus (GEO) under accession number GSE112509, including human nevi (n=23) and melanoma (n=57) samples. Bulk RNA-Seq analysis was performed as described previously (1). Differential mRNA gene expression was determined and normalized with the DESeq2 tool (Love et al., 2014) (Figure 1A, 1B). To analyze patient overall survival in reference to high or low CXCR2 expression, the Cancer Genome Atlas (TCGA) skin cutaneous melanoma (SKCM) dataset was analyzed using the Gene Expression Profiling Interactive Analysis (GEPIA) program. To generate the overall survival plot, the upper and lower quartiles of *CXCR2* expression were used to stratify the groups (Figure 1C). For analysis of the CXCR2 expression level predictive value in therapeutic response to immune checkpoint blockade in melanoma patients a Kaplan-Meier plot of overall survival was constructed by Tumor Immune Dysfunction and Exclusion (TIDE, http://tide.dfci.harvard.edu/) platform in patients with melanoma (2) stratified by CXCR2 z-scores of RNA-Seq normalized count. The P value was calculated by testing the association between prediction scores and overall survival with the two-sided Wald test in a Cox-PH regression (3) (Figure 1D, E).

**Cell lines and establishment of Inducible melanoma mouse models.** `

B16F0 (Catalog number: CRL-6322) and B16F10 (Catalog number: CCL-6475, ATCC) are spontaneous murine melanoma cell lines derived from a C57BL/6J mouse**.** They are tumorigenic and metastatic clones, respectively. Melan-A is a nontumorigenic murine cell line that is syngeneic with B16 melanoma lines (Bennett et al., 1987). Normal human epidermal melanocytes (NHEMs) were purchased from Lonza (#CC-2504) and cultured in melanocyte growth medium including the MM-4 Bullet kit (Lonza, #CC-2349). To examine the regulation of CXCL1/CXCR2 signaling in NHEM, these cells were stimulated with either vehicle, CXCL1 (100ng/ml) or CXCL1+SX692 (5μM) for 24 hours. MelanA and B16 cells were cultured in DMEM/F12 medium (Gibco, #11330-032) containing 10% FBS (Sigma, PAA, A11-201). After 5 days of culture, the culture medium was removed, cells were trypsinized, aspirated, and collected by centrifugation, resuspended in serum containing medium, and the cell number was determined by Countess II (Invitrogen, C10228). Simultaneously, aliquots of cells were permeabilized and stained with Pacific Blue-KI67 and analyzed by flow cytometry. Experiments were conducted in duplicate and repeated once. Data were statistically analyzed with the Student's t-test. In other experiments, NHEMs were treated with CXCL1 (100ng/ml) or vehicle, or with SX-682 and CXCL1 for 24 hours before RNA was extracted and processed for RNAseq analysis. All cell lines are tested monthly for mycoplasma using a PCR detection system.

**Mouse studies with the CXCR1/CXCR2 Inhibitor, SX-682**

To evaluate the effect of the dual CXCR1/CXCR2 inhibitor SX-682 on tumor growth, $Braf^{V600E}/PTEN^{-/-}$ mice were fed with either SX-682 chow (0.756 g/kg of SX-682, Syntrix Pharmaceuticals, Inc) or control chow for two weeks prior to tumor induction via topical administration of 2μl of 5mM 4-HT onto the dorsal skin of 4-week-old mice for three successive days. The mice continued to be fed on the SX-682 chow or control chow until tumors grew to a volume of 1.5 $cm^3$ or required euthanasia due to ulceration or another humane endpoint. *NRas*$^{Q61R}$/*Ink4a*$^{-/-}$ mice pups exposed to 4-HT and UV irradiation were weaned at 21 days and placed on SX-682 or control chow. Tumor volume was measured by microcalipers and tumor number for each mouse was counted after a period of 5 months.

**Flow cytometry analysis and antibodies**

For flow cytometry analyses, tissues were minced on a programmable dissociator and digested with an enzyme solution of collagenase 1 (1,500 CDU, CAT#234153, Calbiochem), dispase II (1 mg/mL, CAT#13689500, Roche), and DNase 1 (0.1 mg/mL, CAT#260913, Calbiochem). The details of staining and flow cytometry analyses protocols can be found in our previous published methodology (4). Mouse tumors were cut into 1- to 2-mm slices and digested in buffer containing 2 mg/mL collagenase and 0.1 mg/mL DNase I. Digested tumors were passed through a 70-mm strainer to obtain a single-cell suspension. Mouse spleens were pressed through 40 mm strainer using syringe plunger to obtain a single-cell suspension. Red blood cells present in whole blood or cell preparations were removed using ACK Lysing Buffer (CAT#RGF-3015, KD Medical) prior to the staining. Cells were incubated with Ghost Dye TM Violet 510 (Tonbo Biosciences), an amine reactive viability dye used to discriminate live/dead cells and washed with FACS buffer (PBS containing 2% v/v FBS). After blocking Fc receptors with anti-mouse CD16/CD32 mAb in FACS buffer for 15 minutes, cells were incubated with mAbs (BioLegend) to mouse CD45-APC/Cy7, CD3-Alexa Fluor 594, CD4-FITC, CD8a-PE, CD44-APC, CD62L-Alexa Fluor 700, CD25-Percp/Cy5.5, CD69-pacific blue; CD45-Alexa Fluor 488, CD11b-PE, F4/80-Brilliant violet 421, Ly6C-APC, Ly6G-PE/Cy7, MHC II-Alexa

Fluor 700, CD206-Percep/Cy5.5, etc. 1 hour on ice. Cells were washed twice in FACS buffer and data acquired with FACSCanto II (Becton Dickenson). For intracellular staining after surface staining, cells were fixing/permeabilizing using Transcription Factor Buffer Set (Cat:562674, BD Pharmingen) per manufacture's protocol. FACS data, comparing leukocytes from $CXCR2^{-/-}$ to $CXCR2^{WT}$ were analyzed using FlowJo software (Version 10.1).

#### Cytokine Array

Tumor lysate or supernatant of cultured cells were prepared as previously described (4). For cultured cell lines, $3x10^5$ cells were plated per well into 6-well plates and incubated overnight. Cells were then treated for 24 hours with 5μM SX-682 (Syntrix) or DMSO prior to supernatant collection. Supernatants were centrifuged to remove cell debris prior to analysis, and both tumor lysates and cell supernatant were subjected to analysis with the Raybio Mouse Cytokine Antibody Array G-Series 3 (Cat# AAM-CYT-G3-8, RayBiotech) per manufacturer's protocol. The glass chip was scanned on the Cy3 channel of a GenoPix 4000B scanner (Genopix 6.1, Molecular Devices, Sunnyvale, CA). For each spot, the net density was determined by subtracting the background. The relative fold difference in cytokine amount was determined in reference to the amount present on the control samples.

#### RNA extraction and RNAseq analysis

RNA was extracted from tumor tissues or cultured cells using the RNeasy Plus Mini Kit (Cat#74134, Qiagen) per manufacturer's protocol. RNAseq was performed using an

Illumina next-generation sequencing platform at the Vanderbilt University Medical Center VANTAGE (Vanderbilt Technologies for Advanced Genomics) Core Facility. Adapters were trimmed by Cutadapt. After trimming, reads were mapped to the mouse genome GRCm38.p6 using STAR and quantified by featureCounts. DESeq2 was used to normalize expression prior to analysis unless otherwise stated. Data were analyzed using GSEA (4.1.0). Gene sets analyzed include: GOBP_CD4_POSITIVE_ALPHA_ BETA_T_CELL_ACTIVATION, GOBP_LYMPHOCYTE_ACTIVATION, GOBP_ LEUKOCYTE_PROLIFERATION, GOBP_LYMPHOCYTE_ANERGY, GOBP_IMMUNE_ RESPONSE, GOBP_STEM_CELL_DIFFERENTIATION, GOBP_ADAPTIVE_IMMUNE_ RESPONSE, GOBP_STEM_CELL_DIVISION, GOBP_CD8_POSITIVE_ALPHA_BETA_ T_CELL_ACTIVATION, GOBP_T_CELL_MEDIATED_IMMUNE_RESPONSE_TO_TUMOR_CELL, GOBP_IMMUNE_RESPONSE_TO_TUMOR_CELL, GOBP_ANTIGEN_ PROCESSING_ AND_ PRESENTATION, GOBP_CD8_POSITIVE_ALPHA_BETA_T_ CELL_PROLIFERATION, GOBP_T_CELL_MEDIATED_CYTOTOXICITY, GOBP_ CELL_CYCLE_PROCESS, GOBP_CELL_CYCLE, GOBP_CELL_CYCLE_G1_S_ PHASE_TRANSITION, and GOBP_MELANOCYTE_PROLIFERATION (Human Molecular Signatures Database-MSigDB). For GSEA, 1000 permutations were performed and human orthologs were annotated with Mouse_Gene_Symbol_Remapping_Human_ Orthologs_MSigDB.v2022.1.Hs.chip. Phenotype permutation was used except when the sample size per phenotype was less than 7. In this case, we follow the GSEA recommendation to use gene set permutation.

For expression plots, DESeq2 normalized counts were transformed by adding one to the value (to retain 0s) and then taking the natural log to account for variance. Comparisons were made using a Mann-Whitney test where appropriate.

For murine microenvironment cell population counter (mMCPcounter) analysis, each count file of protein coding genes was normalized to transcripts per million (TPM) as recommended by the program. TPM was calculated by determining rate as the raw counts divided by the gene length, and then dividing each rate by the sum of all rates multiplied by 1e6. Following TPM normalization, data were transformed utilizing log2(TPM normalized expression +1). The mMCPcounter (v1.1.0) package was installed and the function mMCPcounter.estimate was used to determine predicted immune cell infiltrate.

**Weighted gene co-expression network analysis (WGCNA)**

WGCNA (5) was performed on log-transformed, normalized RNA-seq data of 16382 genes from 15 mouse tumor samples (n = 7 WT mice and n = 8 KO mice) using the R package “WGCNA.” A signed network was generated, such that only positively correlated genes were grouped into modules (negative correlations are given a score of 0), as follows. The function pickSoftThreshold was used to determine the exponent of the correlation coefficients matrix which, when used as weights of network connections, best produces a scale-free network. A power value of 20 was chosen. A topological overlap matrix of network adjacencies between genes was then generated with the adjacency and TOMsimilarity functions, using the Pearson correlation function, which gave a distance measure to be used with average-linkage hierarchical clustering. The WGCNA function cutTreeDynamic was used, with minimum module size of 100, deepSplit = 2, and pamRespectsDendro = FALSE, to generate modules of co-expressed genes. Differentially expressed gene modules across WT and KO conditions were determined using an ANOVA statistical test with FDR correction using the python function selectFDR from the Sci-Kit Learn package. Six gene modules could significantly distinguish between the WT and KO tumors (FDR-adjusted p-value < 0.05) (6).

**Identification of Transcription Factor Network Structure**

To determine the most important transcription factors (TFs) that both distinguish WT and KO conditions and were central to a gene module, we calculated the module membership value, or kME, for each TF, a measure of correlation of its gene expression with the module eigengene from WGCNA. TFs were filtered to those with a significant FDR-adjusted p-value for the ANOVA between WT and KO (< 0.05). We then chose the top 40 significant TFs by kME for each of the six significant gene modules to find the most central TFs for each module.

**Epigenome mapping by ChIPmentation for TCPF21**

ChIPmentation was performed as previously described (7,8) with some modifications. Cells were treated with 5μM SX-682 or DMSO vehicle for 24 hours, collected by trypsinization, and resuspended in PBS. Next, cells were fixed with fresh formaldehyde at final concentration of 1% for 10 min at room temperature. Glycine was added to final concentration of 0.125 M to quench the reaction. Cells were resuspended at $1x10^7$/ml in sonication buffer (10 mM Tris-HCl pH 8.0, 2 mM EDTA pH 8.0, 1% SDS, with Protease Inhibitor Cocktail (Roche) and PhosSTOP™ Phosphatase Inhibitor Cocktail (Roche) and sonicated for 25-30 sec with Diagenode One sonicator in a 50 ul Bioruptor® One Microfluidic Chip till most DNA fragments were in the range of 200 to 700 bp. After sonication, the lysate was adjusted to RIPA buffer conditions (10 mM Tris-HCl pH 8.0, 1 mM EDTA pH 8.0, 140 mM NaCl, 1% Triton X-100, 0.1% SDS, 0.1% sodium deoxycholate, with Protease Inhibitor Cocktail and PhosSTOP™ Phosphatase Inhibitor Cocktail). For each immunoprecipitation, lysate from $2x10^6$ cells and 6 µg of anti-TFCP2L1 antibody (Boster Bio) were used. 6.0 µg of normal rabbit IgG was used as a control. 40µl Dynabeads ™ Protein A (Invitrogen) were used in each immunoprecipitation. After immunoprecipitation, beads were washed twice with RIPA low-salt buffer, twice with RIPA high-salt buffer, twice RIPA lithium-chloride buffer, and once with 10 mM Tris-HCl buffer (pH 8.0). Illumina sequencing adapters were added on bead-bound DNA fragments via tagmetation by using Illumina Tagment DNA TDE1 Enzyme and Buffer Kits (Illumina). After tagmentation, bead-bound DNA fragments were extracted by reversing the crosslink and proteinase K digestion. DNA fragments were purified with AMPure XP beads (Beckman Coulter). qPCR was performed first with KAPA HiFi HotStart Ready Mix and a pair of Nextera custom primers to determine the optimum number of PCR cycles for the DNA library preparation for each immunoprecipitation. The final enriched DNA library for each immunoprecipitation was created by PCR with KAPA HiFi HotStart Ready Mix and Nextera custom primers using the optimum number of PCR cycles determined by qPCR. The enriched DNA library was purified with AMPure XP beads, and size-selection was performed by controlling the concentration of ethanol in the AMPure XP beads and DNA mixture.  Acquired DNA libraries were sequenced by the Illumina NovaSeq 6000 platform at Vanderbilt Technologies for Advanced Genomics core laboratory. ChIPmentation reads were aligned to the mouse reference genome mm10 using Bowtie2 (9). Peaks for each sample were called by MACS2 with an FDR cutoff of 0.01 and the corresponding

IgG input as the control (10-12). Peaks were annotated using Homer (http://homer.ucsd.edu/homer/) and assigned to their closest genes. Enriched motifs were identified by the Homer command findMotifsGenome with the default region size and the motif length (-size 200 and -len 8, 10, 12). Genes that had TFCP2L1 bound to their 5' promoter in the DMSO and SX-682-treated samples were selected and analyzed further using Metascape software (13). Comparisons were made among RPPA, RNAseq, and ChIPseq data sets for B16F0 cells using Metascape software to identify commonly enriched pathways.

**RT-qPCR**

B16F0, B16F10, and MelanA cells were treated with 5µM SX-682 or DMSO for 12 hours prior to collection and RNA extraction with Trizol. cDNA was generated using a Reverse Transcription kit (Catalog number M510A, Promega), and qPCR was performed using a BioRad CFX-qPCR instrument and SsoAdvanced Universal SYBR Green Supermix (Cat: 172-5270, BioRad). Fold changes were calculated using the formula $2^{-(\Delta\Delta Ct)}$, where $\Delta\Delta Ct = \Delta Ct^{(SX\text{-}682)} - \Delta Ct^{(DMSO)}$. The Ct is the cycle at which the threshold line is crossed. Primers for β-actin, sense-ACATGGCATCATCACCAACTG and antisense-AGAATCCAACACGATGCCGG. TFCP2L1, sense-ACACTACAACCAGCACAACTC and antisense-TGGTACTCTGTG TACTGCAGC. Sox10, sense-ACCTATCAGAGGTGGAGCTGAG and antisense-TGC TGTTCCTTCTTGACCTTG. MITF, sense-AGAGCAGCAGTTCTGCAGAGC and antisense-ATGGCTGGTGTCTGACTCACG. Nanog, sense-TCTGCTACTGAGATG CTCTGC and antisense-ACAGTCCGCATCTTCTGCTTC. FoxD3, sense-AGCGATA

TGTCCGGCCAGACG and antisense-TGAGACTGGCCGTGATGAGCG. Notch1, sense-TGACTGTACTGAGAGCTCCTG and antisense-AAGTACCATAGCTGTCTTGGC. TLF4, sense-ATGGCTGTCAGCGACGCTCTG and antisense-TGTTACTGCTGCAAGCTG CAC. Slc4A1, sense-ACTGGAGAACATAATAGGACAG and antisense-AAGGTCAGG TAAGATAGATGTG. Esrrb, sense-AGTGCGAGTATATGCTTAATG and antisense-TGAATTGTCCTCTTGAAGAAG. Sox2, sense-ATGATGGAGACGGAGCTGAAG and antisense-TTGCTGATCTCCGAGTTGTGC. HMGA2, sense-TGCCACAGAAGCGAGGA CGCG and antisense-TCCTAGGTCTGCCTCTTGGCC.

**Protein extraction and RPPA analysis**

Melan-A, B16F0, and B16F10 cells were seeded at a density of $3x10^5$ per well in a 6-well plate overnight prior to 24-hour treatment with 5μM SX-682. Cells were then lysed in a buffer containing 1% Triton X-100, 50mM HEPES, pH 7.4, 150mM NaCl, 1mM EGTA, 100mM NaF, 1.5mM $MgCl_2$, 10mM Na pyrophosphate, 1mM $Na_3VO_4$, 10% glycerol, and freshly added protease inhibitors (Cat# 05056489001, Roche) and phosphatase inhibitors (Cat# 04906837001, Roche) in 2mL reinforced tubes (Cat# P000943-LYSKO-A, Precellys). The homogenization sets at 5000 and goes for 10 sec for cells and 30 sec for tissue in the Precellys 24 Tissue Homogenizer. Lysates were centrifuged at 14,000rpm, 4°C for 2 minutes, and supernatants were collected. Protein concentration was adjusted to 1.5mg/mL and samples were stored in sample buffer (40% glycerol, 8% SDS, 0.25M Tris-HCL, pH 6.8. 10% β-mercaptoethanol is added just before use.) Samples were stored at -80°C until they were submitted to MD Anderson Cancer Center for processing and RPPA analysis of 480 proteins.

RPPA data were normalized and analyzed after a log2 transformation. All comparisons were performed using Welch's t-test. Changes in protein expression were considered significant based upon two criteria: p-value < 0.05 and a $\log_2$(fold change) of ±2 (equivalent to a fold change of ±4). Functional protein association networks are presented using STRING version 11.5.

**Statistical analysis**

Data were summarized in figures using either the mean ± standard deviation (SD) for error bar charts or median with the first and third quartiles for boxplots. The mean of two groups were compared using Welch's t-test. For a comparison of more than two group means, the one-way analysis of variance (ANOVA) with post-hoc Tukey's test was used. The mean difference between two groups by the other factor was assessed in the context of a two-way ANOVA. Pairwise differences between two groups were compared using model-based mean comparisons. The Benjamini and Hochberg (BH) was used to adjust p-value for multiple comparisons as noted in the text to control the within experiment false discovery rate to less than 5%. Data was analyzed on a natural log scale or rank-based scales to meet the normality assumptions of statistical tests as needed. Survival curves are estimated using the Kaplan-Meier method and compared between two groups using the log-rank test. Levels of statistical significance are denoted * for p<0.05, ** for p<0.01, and *** for p<0.001, respectively. The analyses were performed using R version 4.1.2 or GraphPad Prism software. WGCNA statistical analysis was performed as described in the WGCNA methods.

Figure S1.

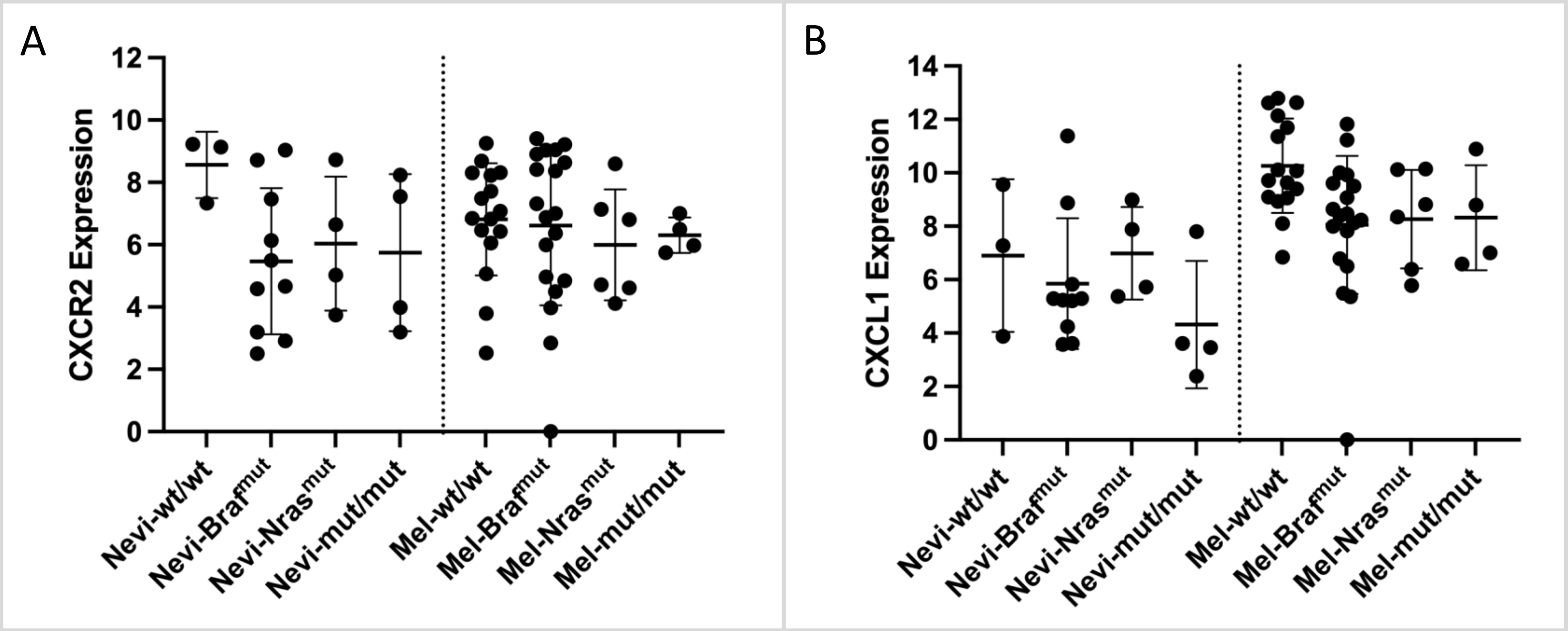
A
CXCR2 Expression
12
10
8
6
4
2
0
Nevi-wt/wt
Nevi-Braf^mut
Nevi-Nras^mut
Nevi-mut/mut
Mel-wt/wt
Mel-Braf^mut
Mel-Nras^mut
Mel-mut/mut
B
CXCL1 Expression
14
12
10
8
6
4
2
0
Nevi-wt/wt
Nevi-Braf^mut
Nevi-Nras^mut
Nevi-mut/mut
Mel-wt/wt
Mel-Braf^mut
Mel-Nras^mut
Mel-mut/mut

Figure S2.

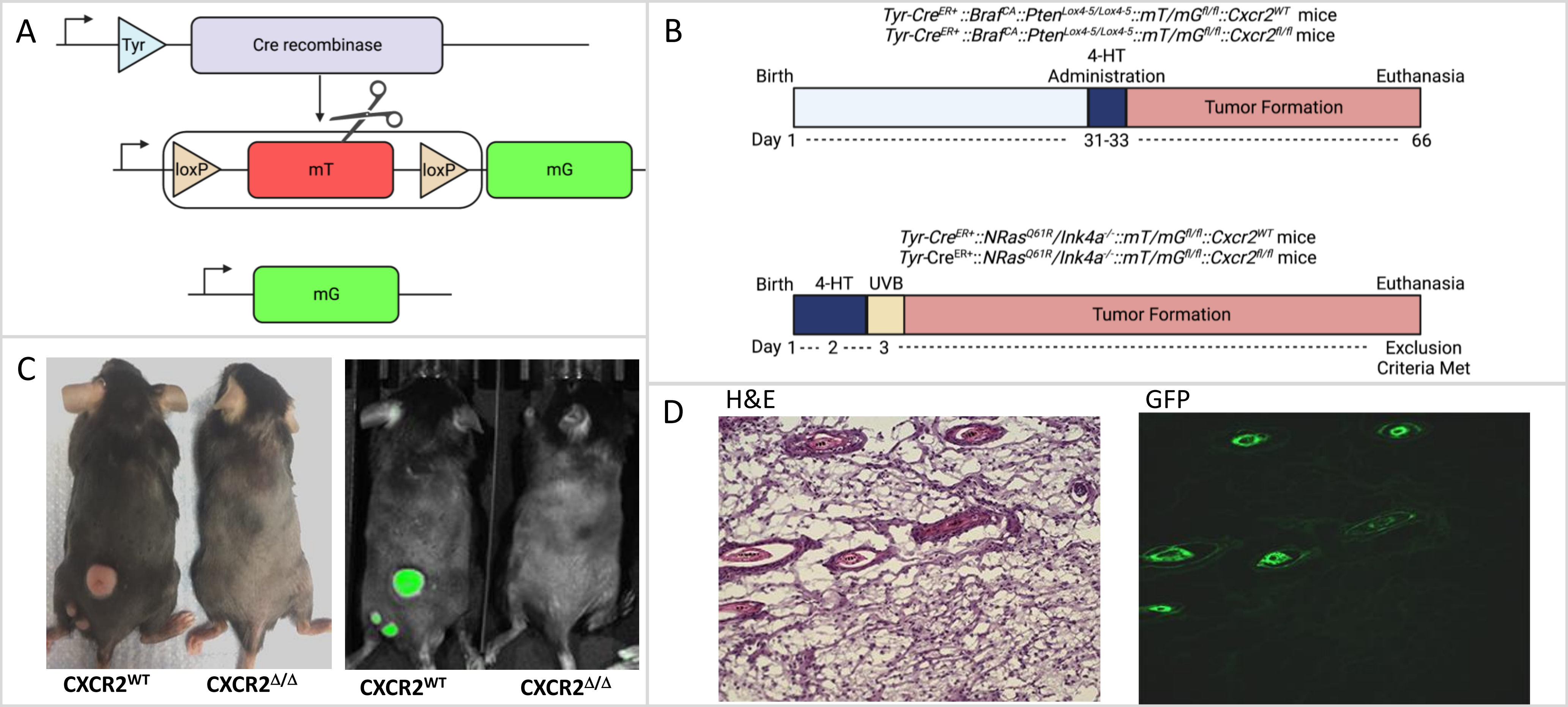
A
Tyr
Cre recombinase
loxP
mT
loxP
mG
mG
B
Tyr-CreER+::BrafCA::PtenLox4-5/Lox4-5::mT/mGfl/fl::Cxcr2WT mice
Tyr-CreER+::BrafCA::PtenLox4-5/Lox4-5::mT/mGfl/fl::Cxcr2fl/fl mice
Birth
4-HT Administration
Euthanasia
Tumor Formation
Day 1
31-33
66
Tyr-CreER+::NRasQ61R/Ink4a-/-::mT/mGfl/fl::Cxcr2WT mice
Tyr-CreER+::NRasQ61R/Ink4a-/-::mT/mGfl/fl::Cxcr2fl/fl mice
Birth
4-HT
UVB
Euthanasia
Tumor Formation
Day 1
2
3
Exclusion Criteria Met
C
CXCR2WT
CXCR2Δ/Δ
CXCR2WT
CXCR2Δ/Δ
D
H&E
GFP

### Figure S3

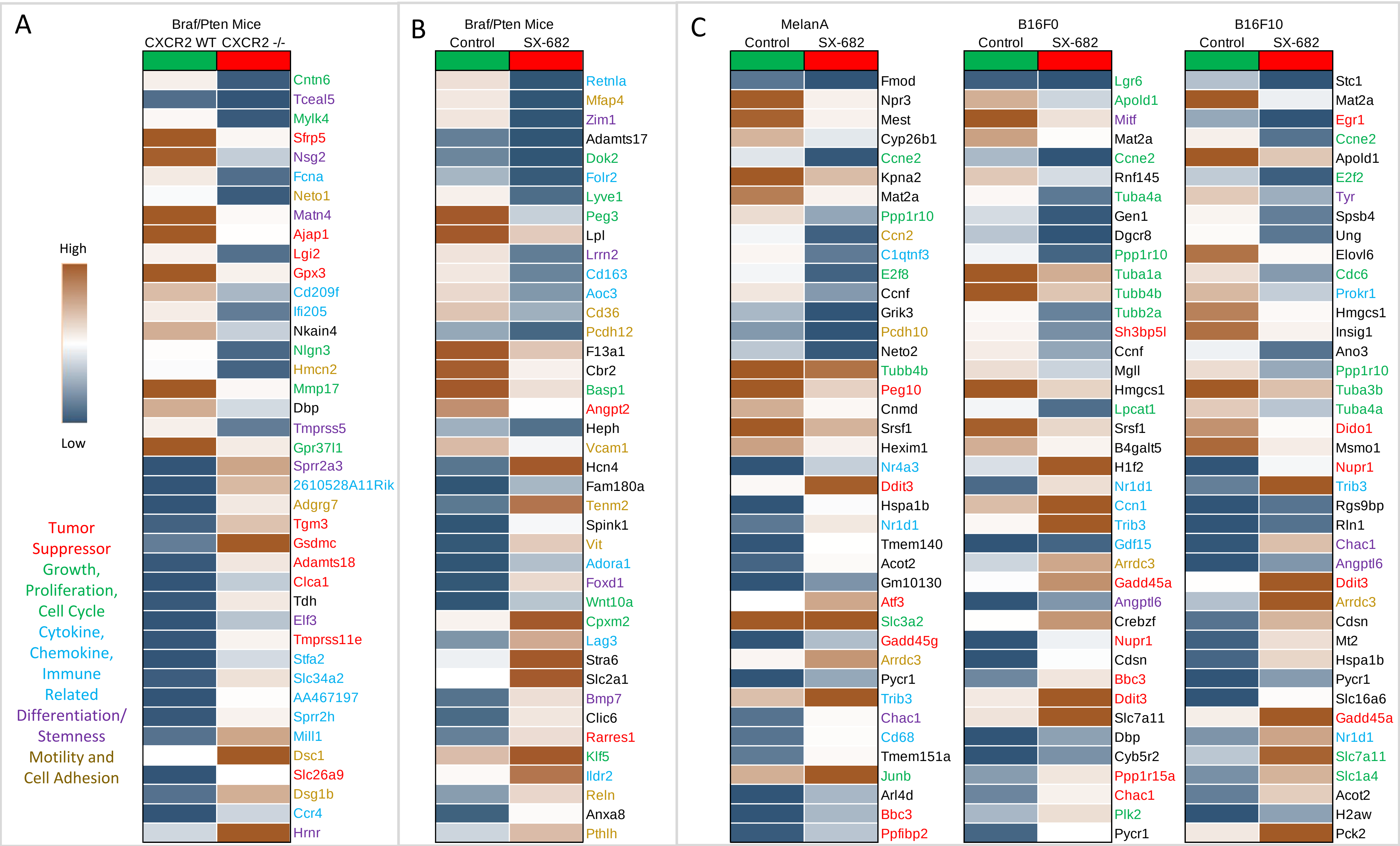
A
Braf/Pten Mice
CXCR2 WT
CXCR2 -/-
High
Low
Tumor Suppressor
Growth, Proliferation, Cell Cycle
Cytokine, Chemokine, Immune Related
Differentiation/ Stemness
Motility and Cell Adhesion
Cntn6
Tceal5
Mylk4
Sfrp5
Nsg2
Fcna
Neto1
Matn4
Ajap1
Lgi2
Gpx3
Cd209f
Ifi205
Nkain4
Nlgn3
Hmcn2
Mmp17
Dbp
Tmprss5
Gpr37l1
Sprr2a3
2610528A11Rik
Adgrg7
Tgm3
Gsdmc
Adamts18
Clca1
Tdh
Elf3
Tmprss11e
Stfa2
Slc34a2
AA467197
Sprr2h
Mill1
Dsc1
Slc26a9
Dsg1b
Ccr4
Hrnr
B
Braf/Pten Mice
Control
SX-682
Retnla
Mfap4
Zim1
Adamts17
Dok2
Folr2
Lyve1
Peg3
Lpl
Lrrn2
Cd163
Aoc3
Cd36
Pcdh12
F13a1
Cbr2
Basp1
Angpt2
Heph
Vcam1
Hcn4
Fam180a
Tenm2
Spink1
Vit
Adora1
Foxd1
Wnt10a
Cpxm2
Lag3
Stra6
Slc2a1
Bmp7
Clic6
Rarres1
Klf5
Ildr2
Reln
Anxa8
Pthlh
C
MelanA
Control
SX-682
Fmod
Npr3
Mest
Cyp26b1
Ccne2
Kpna2
Mat2a
Ppp1r10
Ccn2
C1qtnf3
E2f8
Ccnf
Grik3
Pcdh10
Neto2
Tubb4b
Peg10
Cnmd
Srsf1
Hexim1
Nr4a3
Ddit3
Hspa1b
Nr1d1
Tmem140
Acot2
Gm10130
Atf3
Slc3a2
Gadd45g
Arrdc3
Pycr1
Trib3
Chac1
Cd68
Tmem151a
Junb
Arl4d
Bbc3
Ppfibp2
B16F0
Control
SX-682
Lgr6
Apold1
Mitf
Mat2a
Ccne2
Rnf145
Tuba4a
Gen1
Dgcr8
Ppp1r10
Tuba1a
Tubb4b
Tubb2a
Sh3bp5l
Ccnf
Mgll
Hmgcs1
Lpcat1
Srsf1
B4galt5
H1f2
Nr1d1
Ccn1
Trib3
Gdf15
Arrdc3
Gadd45a
Angptl6
Crebzf
Nupr1
Cdsn
Bbc3
Ddit3
Slc7a11
Dbp
Cyb5r2
Ppp1r15a
Chac1
Plk2
Pycr1
B16F10
Control
SX-682
Stc1
Mat2a
Egr1
Ccne2
Apold1
E2f2
Tyr
Spsb4
Ung
Elovl6
Cdc6
Prokr1
Hmgcs1
Insig1
Ano3
Ppp1r10
Tuba3b
Tuba4a
Dido1
Msmo1
Nupr1
Trib3
Rgs9bp
Rln1
Chac1
Angptl6
Ddit3
Arrdc3
Cdsn
Mt2
Hspa1b
Pycr1
Slc16a6
Gadd45a
Nr1d1
Slc7a11
Slc1a4
Acot2
H2aw
Pck2

Figure S4.

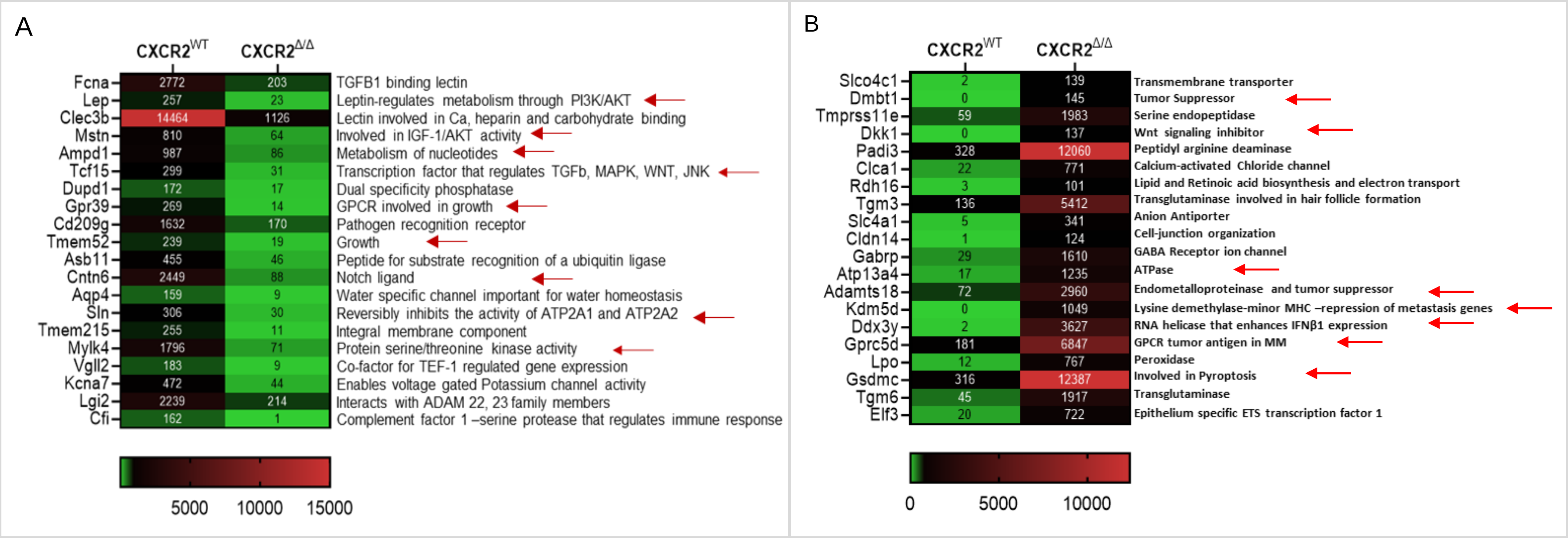
A
CXCR2WT
CXCR2Δ/Δ
Fcna 2772 203 TGFB1 binding lectin
Lep 257 23 Leptin-regulates metabolism through PI3K/AKT
Clec3b 14464 1126 Lectin involved in Ca, heparin and carbohydrate binding
Mstn 810 64 Involved in IGF-1/AKT activity
Ampd1 987 86 Metabolism of nucleotides
Tcf15 299 31 Transcription factor that regulates TGFb, MAPK, WNT, JNK
Dupd1 172 17 Dual specificity phosphatase
Gpr39 269 14 GPCR involved in growth
Cd209g 1632 170 Pathogen recognition receptor
Tmem52 239 19 Growth
Asb11 455 46 Peptide for substrate recognition of a ubiquitin ligase
Cntn6 2449 88 Notch ligand
Aqp4 159 9 Water specific channel important for water homeostasis
Sln 306 30 Reversibly inhibits the activity of ATP2A1 and ATP2A2
Tmem215 255 11 Integral membrane component
Mylk4 1796 71 Protein serine/threonine kinase activity
Vgll2 183 9 Co-factor for TEF-1 regulated gene expression
Kcna7 472 44 Enables voltage gated Potassium channel activity
Lgi2 2239 214 Interacts with ADAM 22, 23 family members
Cfi 162 1 Complement factor 1 –serine protease that regulates immune response
5000 10000 15000
B
CXCR2WT
CXCR2Δ/Δ
Slco4c1 2 139 Transmembrane transporter
Dmbt1 0 145 Tumor Suppressor
Tmprss11e 59 1983 Serine endopeptidase
Dkk1 0 137 Wnt signaling inhibitor
Padi3 328 12060 Peptidyl arginine deaminase
Clca1 22 771 Calcium-activated Chloride channel
Rdh16 3 101 Lipid and Retinoic acid biosynthesis and electron transport
Tgm3 136 5412 Transglutaminase involved in hair follicle formation
Slc4a1 5 341 Anion Antiporter
Cldn14 1 124 Cell-junction organization
Gabrp 29 1610 GABA Receptor ion channel
Atp13a4 17 1235 ATPase
Adamts18 72 2960 Endometalloproteinase and tumor suppressor
Kdm5d 0 1049 Lysine demethylase-minor MHC –repression of metastasis genes
Ddx3y 2 3627 RNA helicase that enhances IFNβ1 expression
Gprc5d 181 6847 GPCR tumor antigen in MM
Lpo 12 767 Peroxidase
Gsdmc 316 12387 Involved in Pyroptosis
Tgm6 45 1917 Transglutaminase
Elf3 20 722 Epithelium specific ETS transcription factor 1
0 5000 10000

Figure S5.

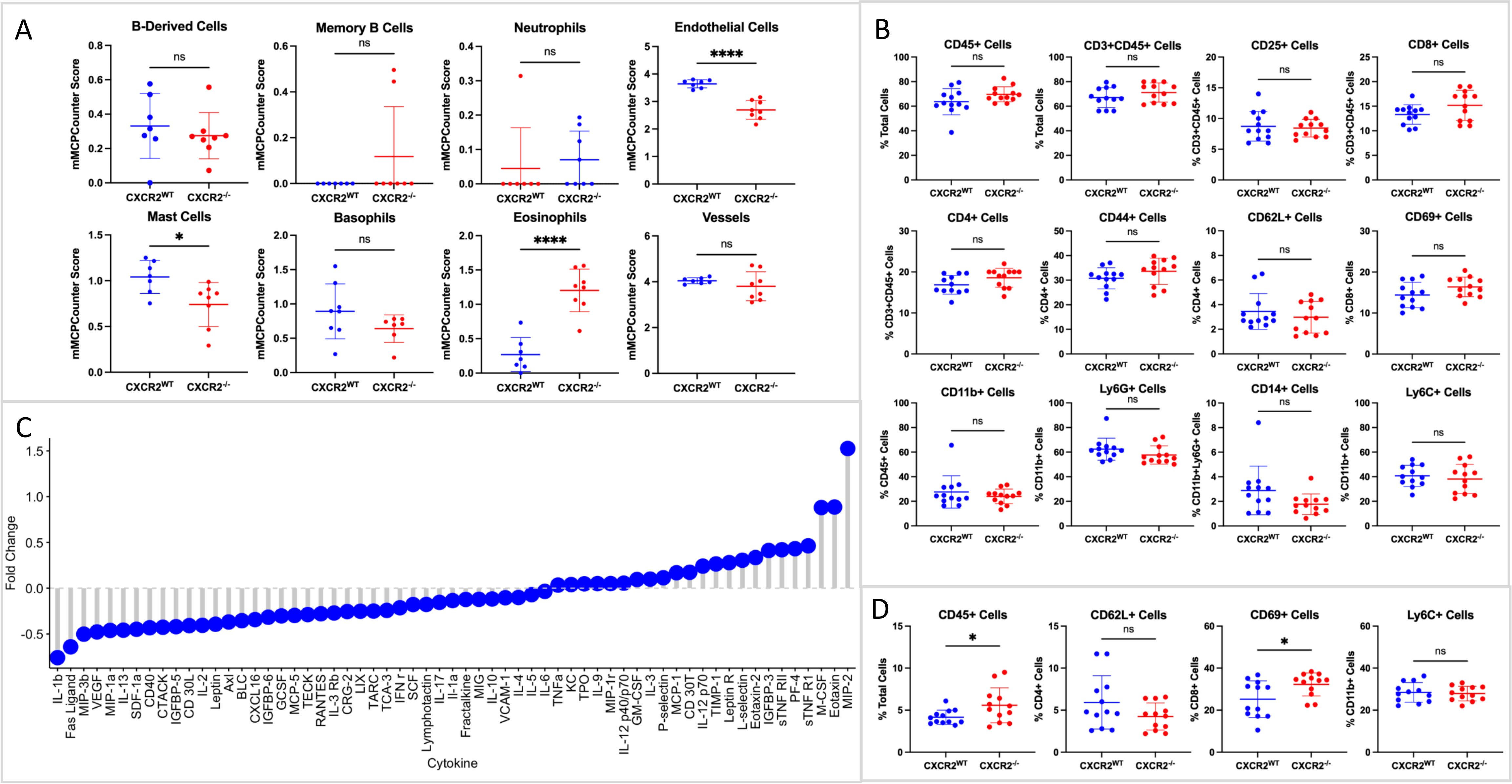
A
B-Derived Cells
Memory B Cells
Neutrophils
Endothelial Cells
Mast Cells
Basophils
Eosinophils
Vessels
mMCPCounter Score
CXCR2WT
CXCR2-/-
ns
*
****
B
CD45+ Cells
CD3+CD45+ Cells
CD25+ Cells
CD8+ Cells
CD4+ Cells
CD44+ Cells
CD62L+ Cells
CD69+ Cells
CD11b+ Cells
Ly6G+ Cells
CD14+ Cells
Ly6C+ Cells
% Total Cells
% CD3+CD45+ Cells
% CD4+ Cells
% CD8+ Cells
% CD45+ Cells
% CD11b+ Cells
% CD11b+Ly6G+ Cells
C
Fold Change
Cytokine
IL-1b
Fas Ligand
MIP-3b
VEGF
MIP-1a
IL-13
SDF-1a
CD40
CTACK
IGFBP-5
CD 30L
IL-2
Leptin
Axl
BLC
CXCL16
IGFBP-6
GCSF
MCP-5
TECK
RANTES
IL-3 Rb
CRG-2
LIX
TARC
TCA-3
IFN r
SCF
Lymphotactin
IL-17
Il-1a
Fractalkine
MIG
IL-10
VCAM-1
IL-4
IL-5
IL-6
TNFa
KC
TPO
IL-9
MIP-1r
IL-12 p40/p70
GM-CSF
IL-3
P-selectin
MCP-1
CD 30T
IL-12 p70
TIMP-1
Leptin R
L-selectin
Eotaxin-2
IGFBP-3
sTNF RII
PF-4
sTNF R1
M-CSF
Eotaxin
MIP-2
D
CD45+ Cells
CD62L+ Cells
CD69+ Cells
Ly6C+ Cells

Figure S6.

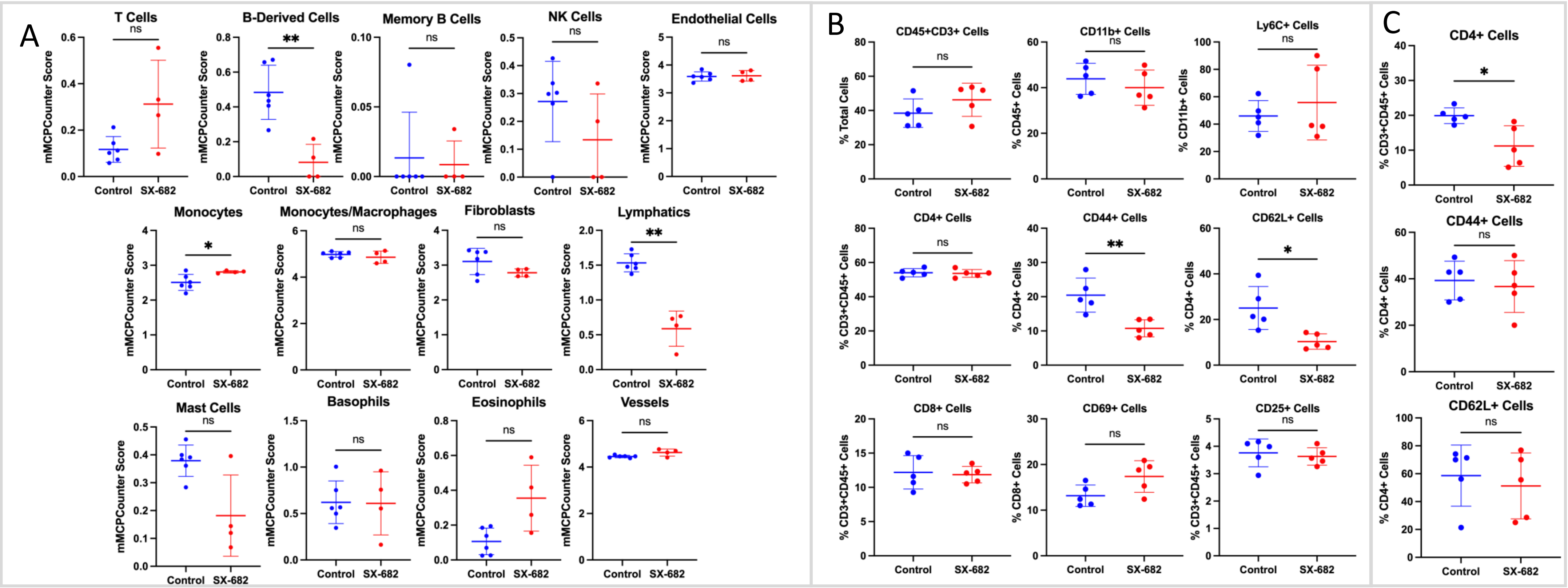

A
T Cells
B-Derived Cells
Memory B Cells
NK Cells
Endothelial Cells
Monocytes
Monocytes/Macrophages
Fibroblasts
Lymphatics
Mast Cells
Basophils
Eosinophils
Vessels
mMCPCounter Score
Control
SX-682
ns
*
**
B
CD45+CD3+ Cells
CD11b+ Cells
Ly6C+ Cells
CD4+ Cells
CD44+ Cells
CD62L+ Cells
CD8+ Cells
CD69+ Cells
CD25+ Cells
% Total Cells
% CD45+ Cells
% CD11b+ Cells
% CD3+CD45+ Cells
% CD4+ Cells
% CD8+ Cells
C
CD4+ Cells
CD44+ Cells
CD62L+ Cells

Figure S7.

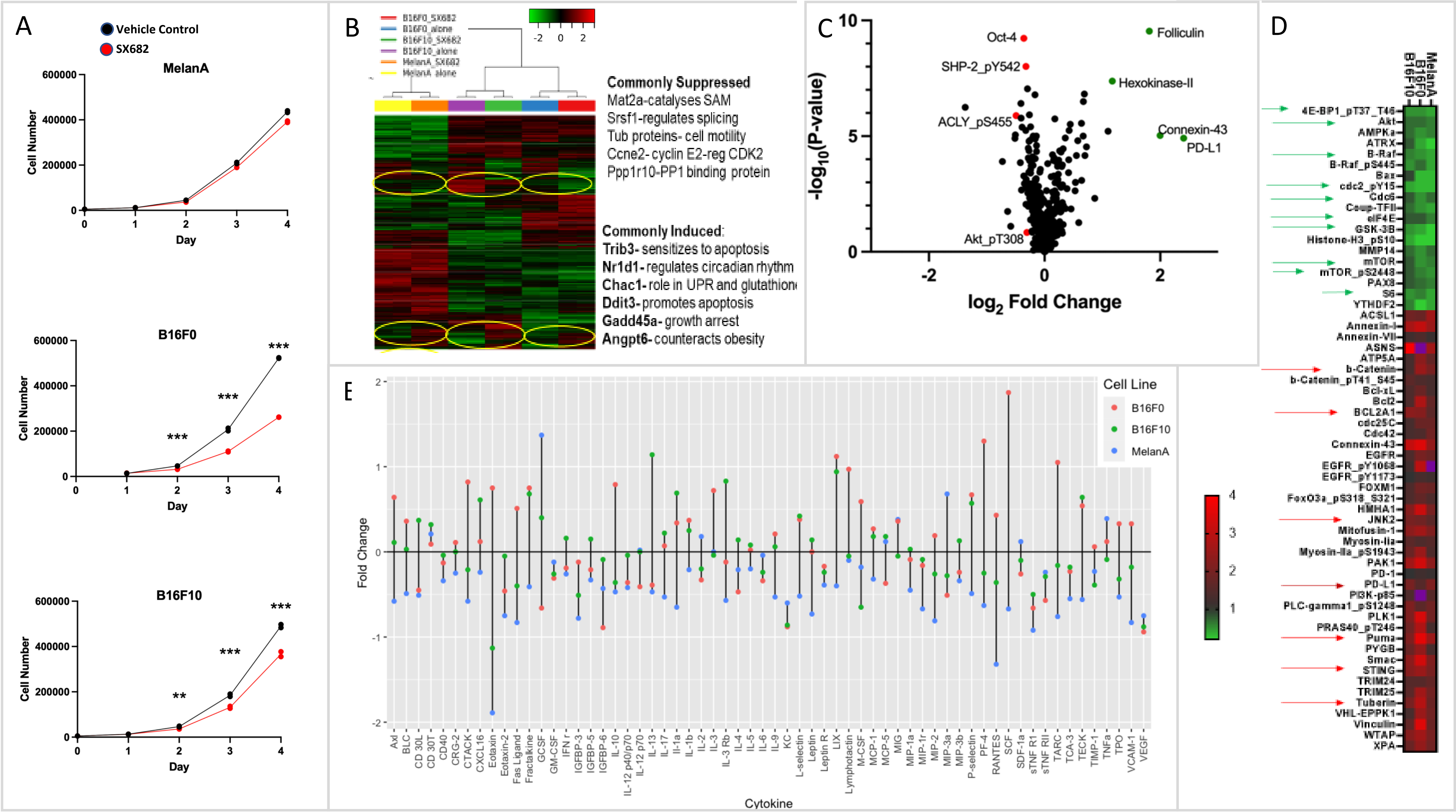
A
Vehicle Control
SX682
MelanA
B16F0
B16F10
Cell Number
Day
B
Commonly Suppressed
Mat2a-catalyses SAM
Srsf1-regulates splicing
Tub proteins- cell motility
Ccne2- cyclin E2-reg CDK2
Ppp1r10-PP1 binding protein
Commonly Induced:
Trib3- sensitizes to apoptosis
Nr1d1- regulates circadian rhythm
Chac1- role in UPR and glutathion
Ddit3- promotes apoptosis
Gadd45a- growth arrest
Angpt6- counteracts obesity
C
Oct-4
SHP-2_pY542
ACLY_pS455
Akt_pT308
Folliculin
Hexokinase-II
Connexin-43
PD-L1
-log10(P-value)
log2 Fold Change
D
MelanA
B16F0
B16F10
E
Cell Line
B16F0
B16F10
MelanA
Fold Change
Cytokine

Figure S8.

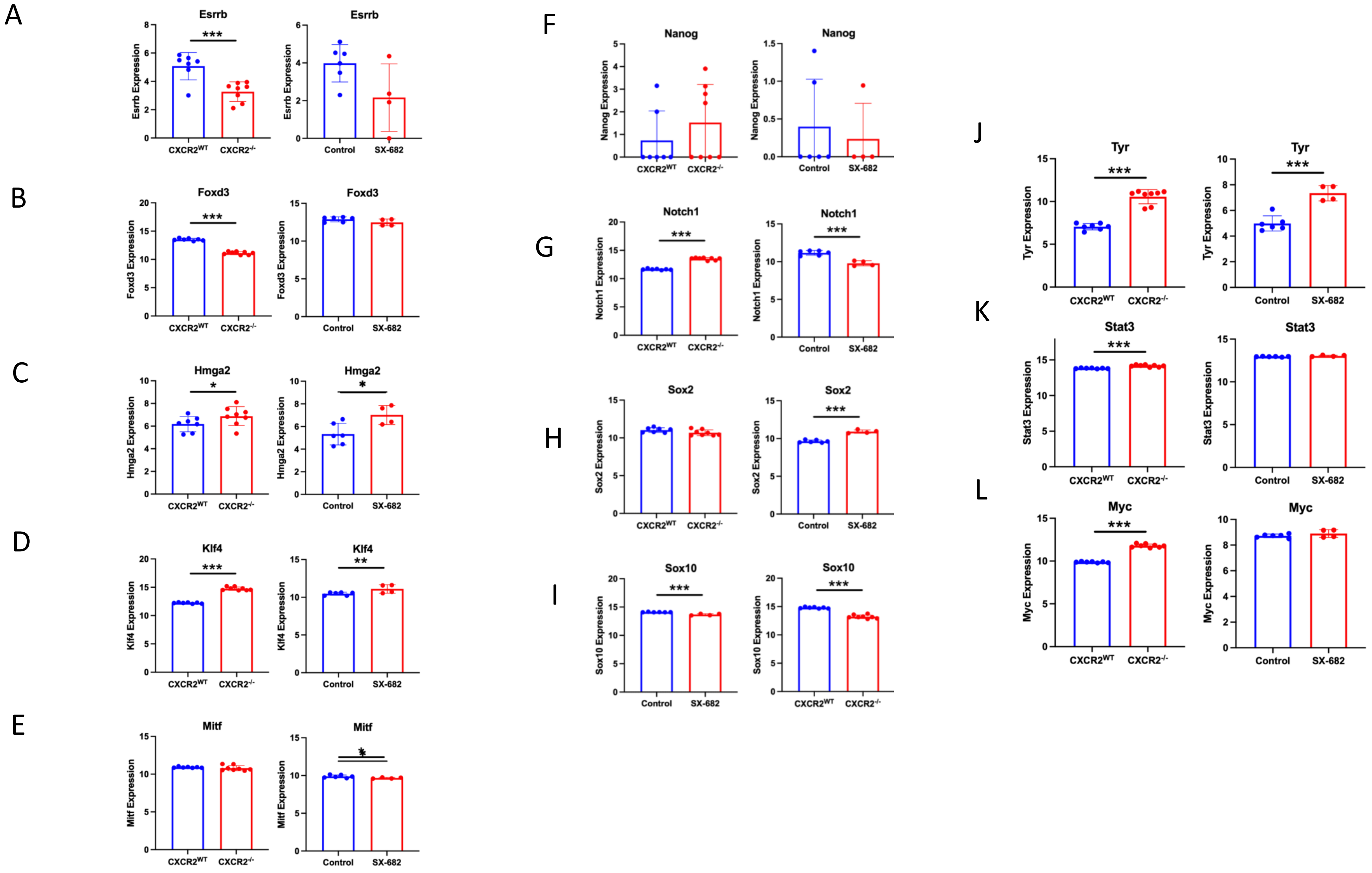

A
Esrrb
***
Esrrb Expression
CXCR2WT
CXCR2-/-
Esrrb
Esrrb Expression
Control
SX-682
B
Foxd3
***
Foxd3 Expression
CXCR2WT
CXCR2-/-
Foxd3
Foxd3 Expression
Control
SX-682
C
Hmga2
*
Hmga2 Expression
CXCR2WT
CXCR2-/-
Hmga2
*
Hmga2 Expression
Control
SX-682
D
Klf4
***
Klf4 Expression
CXCR2WT
CXCR2-/-
Klf4
**
Klf4 Expression
Control
SX-682
E
Mitf
Mitf Expression
CXCR2WT
CXCR2-/-
Mitf
*
Mitf Expression
Control
SX-682
F
Nanog
Nanog Expression
CXCR2WT
CXCR2-/-
Nanog
Nanog Expression
Control
SX-682
G
Notch1
***
Notch1 Expression
CXCR2WT
CXCR2-/-
Notch1
***
Notch1 Expression
Control
SX-682
H
Sox2
Sox2 Expression
CXCR2WT
CXCR2-/-
Sox2
***
Sox2 Expression
Control
SX-682
I
Sox10
***
Sox10 Expression
Control
SX-682
Sox10
***
Sox10 Expression
CXCR2WT
CXCR2-/-
J
Tyr
***
Tyr Expression
CXCR2WT
CXCR2-/-
Tyr
***
Tyr Expression
Control
SX-682
K
Stat3
***
Stat3 Expression
CXCR2WT
CXCR2-/-
Stat3
Stat3 Expression
Control
SX-682
L
Myc
***
Myc Expression
CXCR2WT
CXCR2-/-
Myc
Myc Expression
Control
SX-682

Figure S9

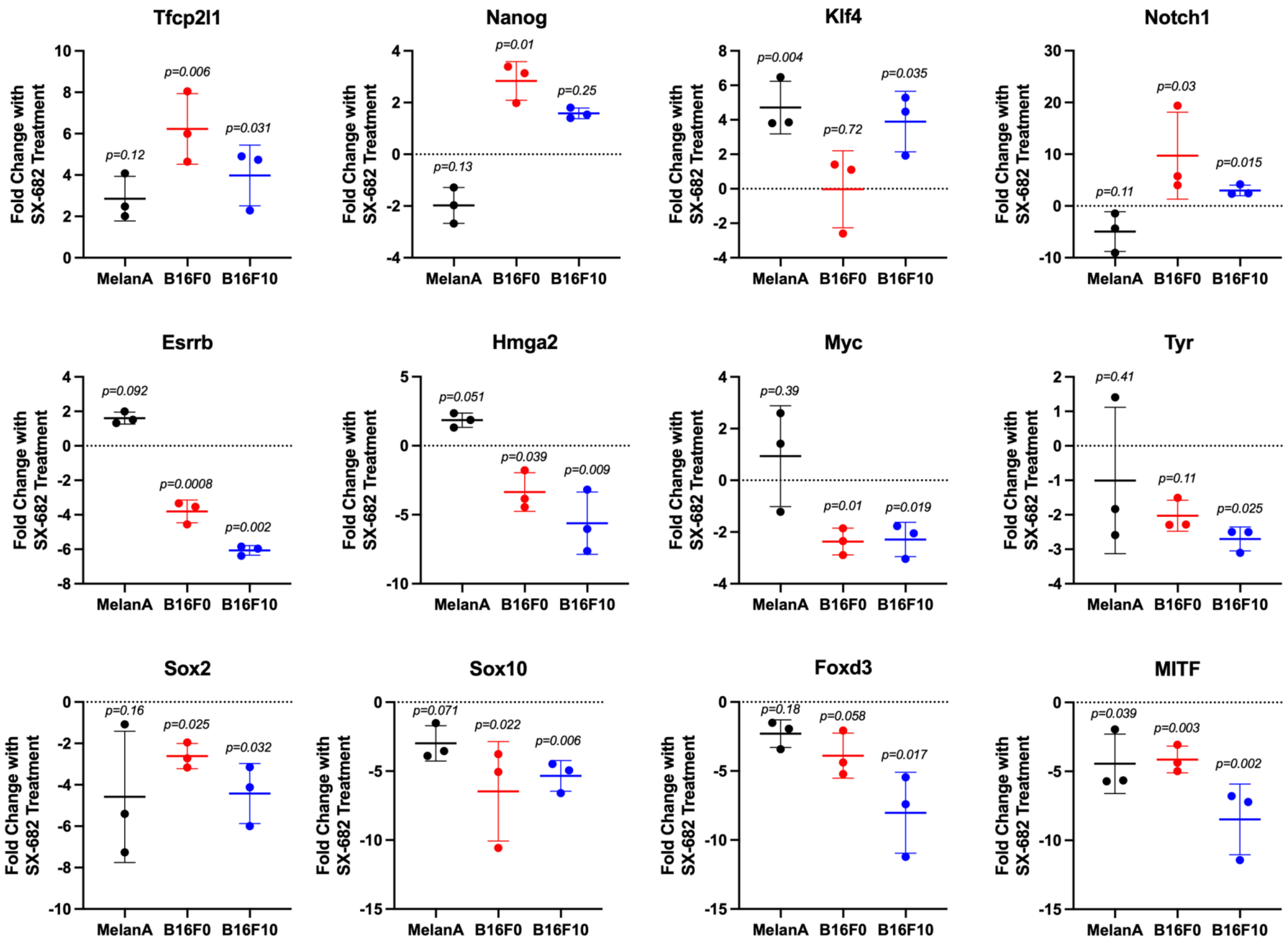

Tfcp2l1
Nanog
Klf4
Notch1
Esrrb
Hmga2
Myc
Tyr
Sox2
Sox10
Foxd3
MITF
Fold Change with SX-682 Treatment
MelanA
B16F0
B16F10
p=0.12
p=0.006
p=0.031
p=0.13
p=0.01
p=0.25
p=0.004
p=0.72
p=0.035
p=0.11
p=0.03
p=0.015
p=0.092
p=0.0008
p=0.002
p=0.051
p=0.039
p=0.009
p=0.39
p=0.01
p=0.019
p=0.41
p=0.11
p=0.025
p=0.16
p=0.025
p=0.032
p=0.071
p=0.022
p=0.006
p=0.18
p=0.058
p=0.017
p=0.039
p=0.003
p=0.002

### Figure S10

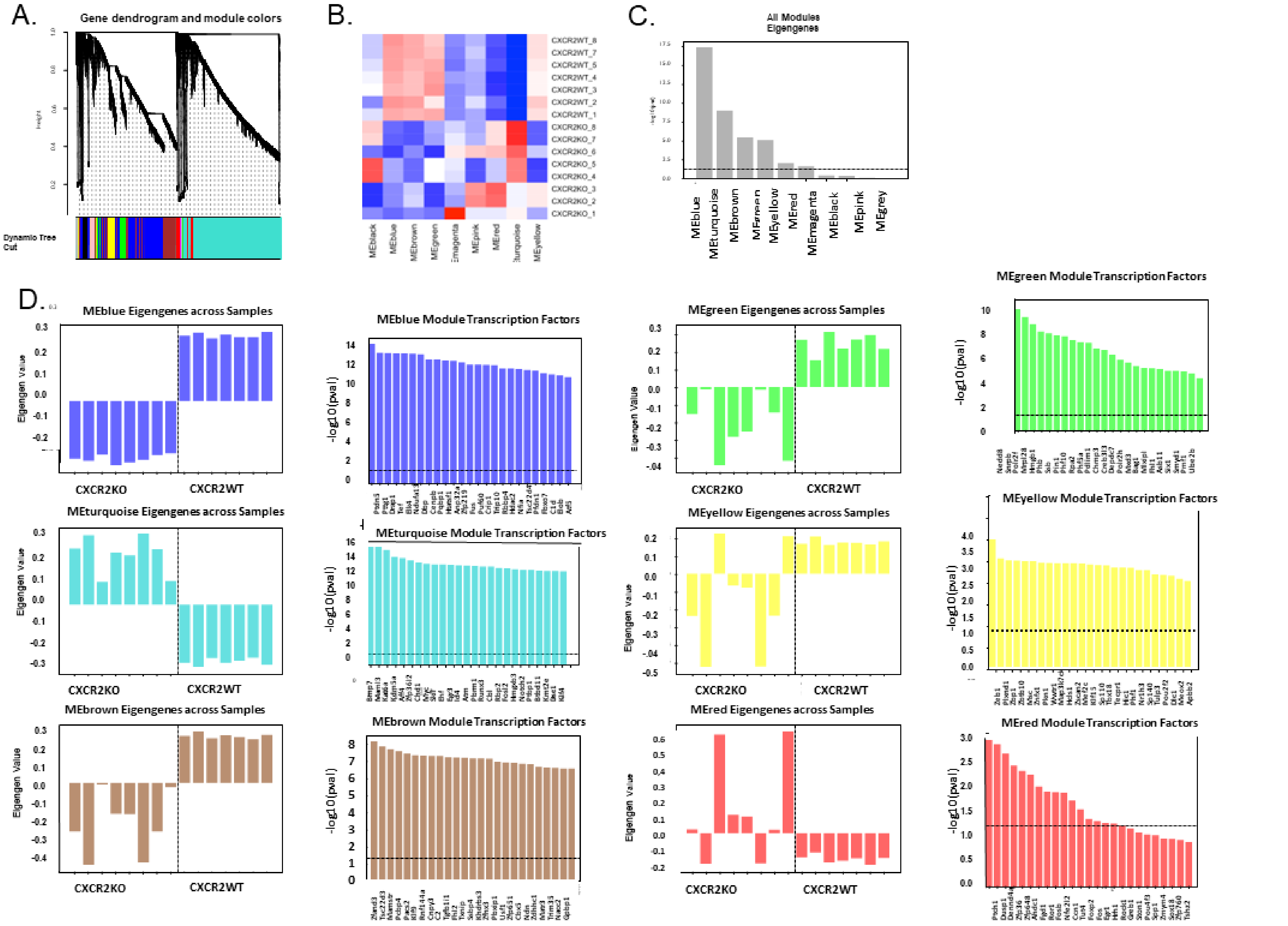
A.
Gene dendrogram and module colors
Dynamic Tree Cut
B.
C.
All Modules Eigengenes
D.
MEblue Eigengenes across Samples
MEblue Module Transcription Factors
MEgreen Eigengenes across Samples
MEgreen Module Transcription Factors
MEturquoise Eigengenes across Samples
MEturquoise Module Transcription Factors
MEyellow Eigengenes across Samples
MEyellow Module Transcription Factors
MEbrown Eigengenes across Samples
MEbrown Module Transcription Factors
MEred Eigengenes across Samples
MEred Module Transcription Factors
Eigengen Value
-log10(pval)
CXCR2KO
CXCR2WT

Figure S11

#### A. Terms from Genes with Increased Expression

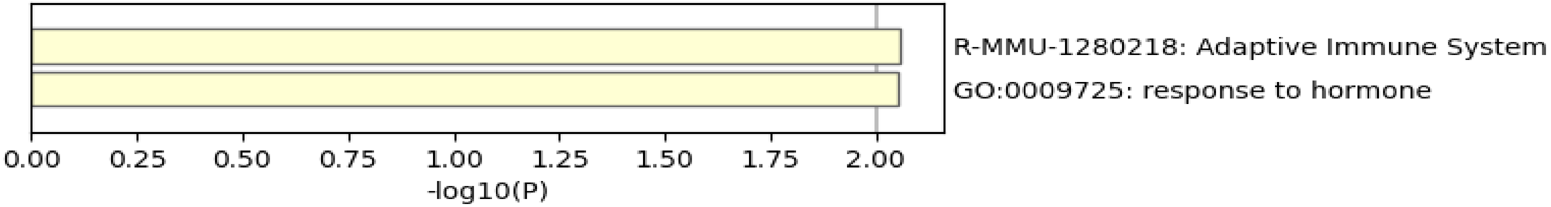


#### B. Terms from Genes with Decreased Expression

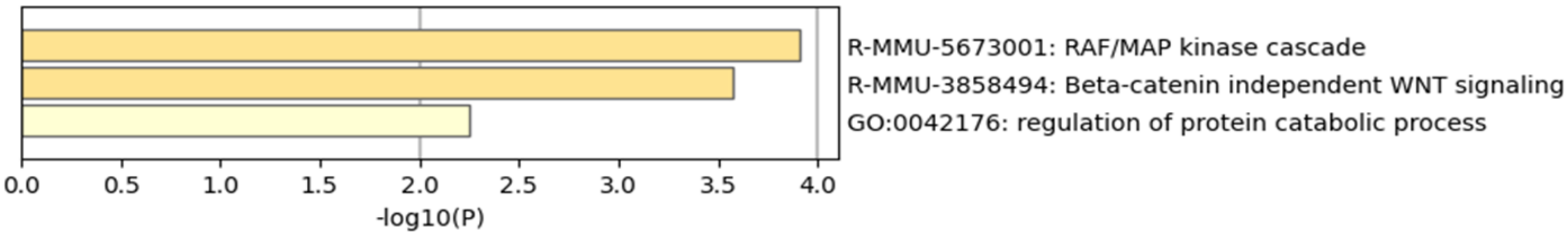


#### .C. Gene Overlap Analysis Heat Map of Selected GO Parent Files

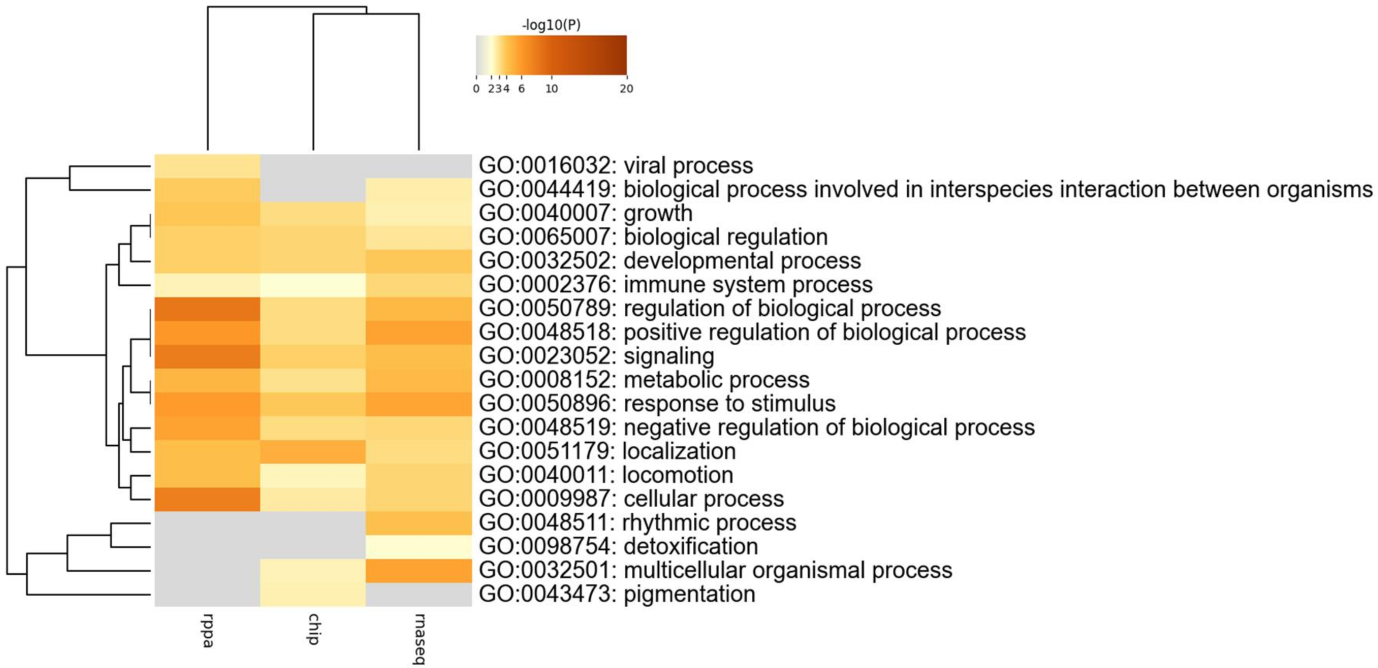


#### Gene Overlap Analysis—Expanded via Shared Enriched Ontologies

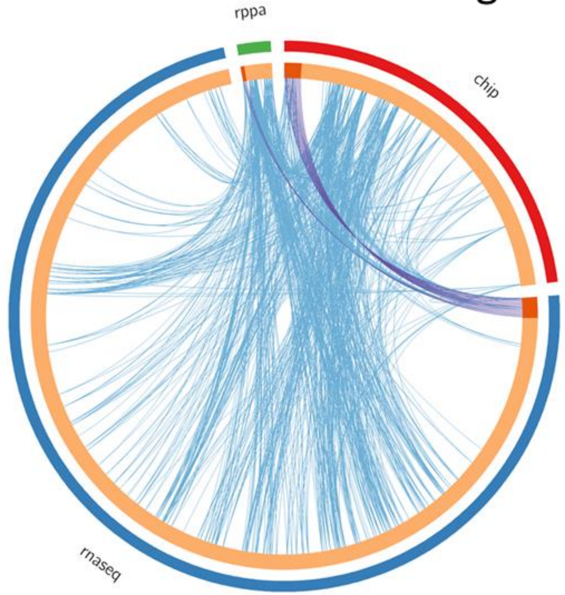

#### Figure S11D

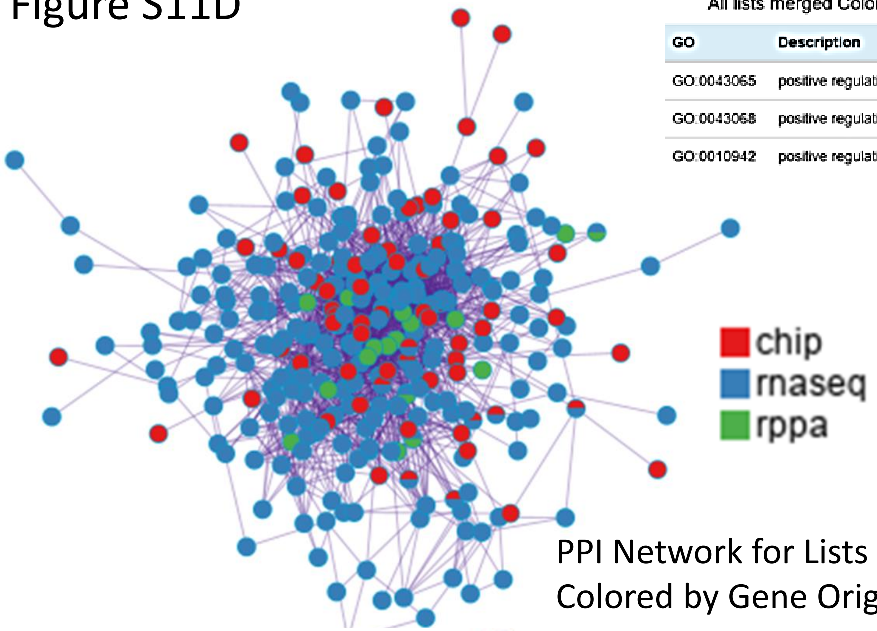


PPI Network for Lists Merged Colored by Gene Origin in Studies

All lists merged Colored by Cluster(Full Connection)

| GO | Description | Log10(P) |
|---|---|---|
| GO:0043065 | positive regulation of apoptotic process | -21.9 |
| GO:0043068 | positive regulation of programmed cell death | -21.6 |
| GO:0010942 | positive regulation of cell death | -19.5 |

#### Figure S11E

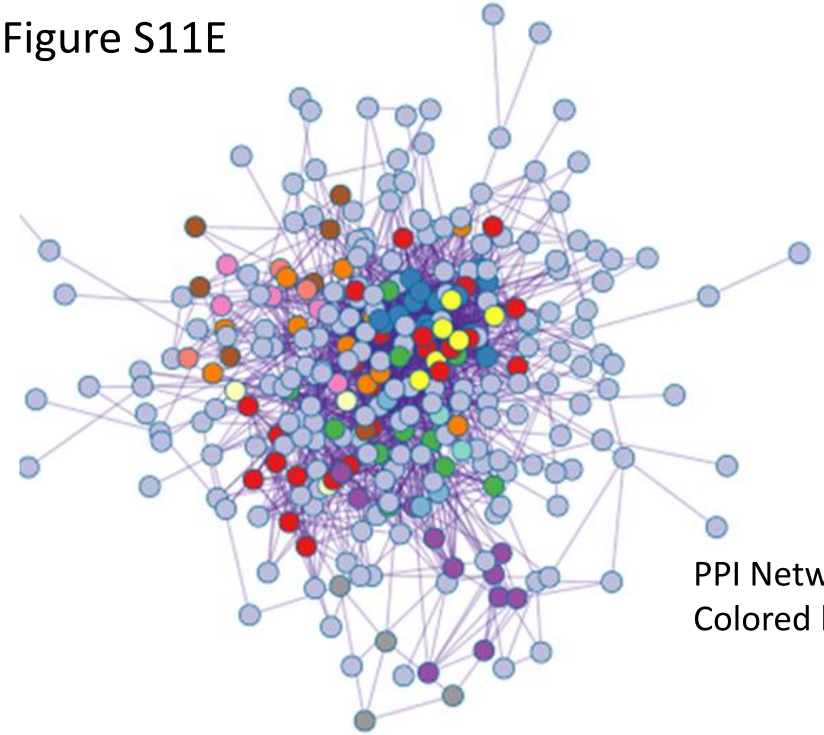

PPI Network for Lists Merged Colored by MCODE ID

All lists merged Colored by Cluster(Keep MCODE Nodes Only)

| Color | MCODE | GO | Description | Log10(P) |
|---|---|---|---|---|
| | MCODE_1 | R-MMU-212300 | PRC2 methylates histones and DNA | -15.6 |
| | MCODE_1 | R-MMU-8936459 | RUNX1 regulates genes involved in megakaryocyte differentiation and platelet function | -15.4 |
| | MCODE_1 | R-MMU-212165 | Epigenetic regulation of gene expression | -14.3 |
| | MCODE_2 | WP6 | Integrin-mediated cell adhesion | -17.3 |
| | MCODE_2 | WP85 | Focal adhesion | -17.1 |
| | MCODE_2 | mmu04510 | Focal adhesion - Mus musculus (house mouse) | -14.5 |
| | MCODE_3 | GO:0006366 | transcription by RNA polymerase II | -6.4 |
| | MCODE_3 | GO:0006351 | DNA-templated transcription | -5.6 |
| | MCODE_3 | GO:0097659 | nucleic acid-templated transcription | -5.6 |
| | MCODE_4 | mmu00480 | Glutathione metabolism - Mus musculus (house mouse) | -12.2 |
| | MCODE_4 | GO:0006749 | glutathione metabolic process | -11.9 |
| | MCODE_4 | R-MMU-156590 | Glutathione conjugation | -11.2 |
| | MCODE_5 | R-MMU-5358351 | Signaling by Hedgehog | -8.3 |
| | MCODE_5 | R-MMU-2262752 | Cellular responses to stress | -7.4 |
| | MCODE_5 | R-MMU-8953897 | Cellular responses to stimuli | -7.4 |
| | MCODE_6 | GO:0007169 | transmembrane receptor protein tyrosine kinase signaling pathway | -7.3 |
| | MCODE_6 | GO:0030335 | positive regulation of cell migration | -6.5 |
| | MCODE_6 | GO:2000147 | positive regulation of cell motility | -6.4 |
| | MCODE_7 | mmu04710 | Circadian rhythm - Mus musculus (house mouse) | -9.7 |
| | MCODE_7 | GO:0032922 | circadian regulation of gene expression | -8.3 |
| | MCODE_7 | GO:0007623 | circadian rhythm | -7.2 |
| | MCODE_8 | R-MMU-72187 | mRNA 3'-end processing | -9.7 |
| | MCODE_8 | R-MMU-73856 | RNA Polymerase II Transcription Termination | -9.4 |
| | MCODE_8 | GO:0006397 | mRNA processing | -8.5 |
| | MCODE_9 | mmu00900 | Terpenoid backbone biosynthesis - Mus musculus (house mouse) | -11.9 |
| | MCODE_9 | WP4346 | Cholesterol metabolism with Bloch and Kandutsch-Russell pathways | -10.5 |
| | MCODE_9 | mmu00650 | Butanoate metabolism - Mus musculus (house mouse) | -8.1 |
| | MCODE_10 | GO:1900745 | positive regulation of p38MAPK cascade | -8.6 |
| | MCODE_10 | mmu05216 | Thyroid cancer - Mus musculus (house mouse) | -8.3 |
| | MCODE_10 | GO:1900744 | regulation of p38MAPK cascade | -8.1 |
| | MCODE_13 | R-MMU-1280218 | Adaptive Immune System | -4.5 |
| | MCODE_14 | GO:0070227 | lymphocyte apoptotic process | -7.7 |
| | MCODE_14 | GO:0071887 | leukocyte apoptotic process | -7.5 |
| | MCODE_14 | GO:0008630 | intrinsic apoptotic signaling pathway in response to DNA damage | -7.2 |

Figure S12

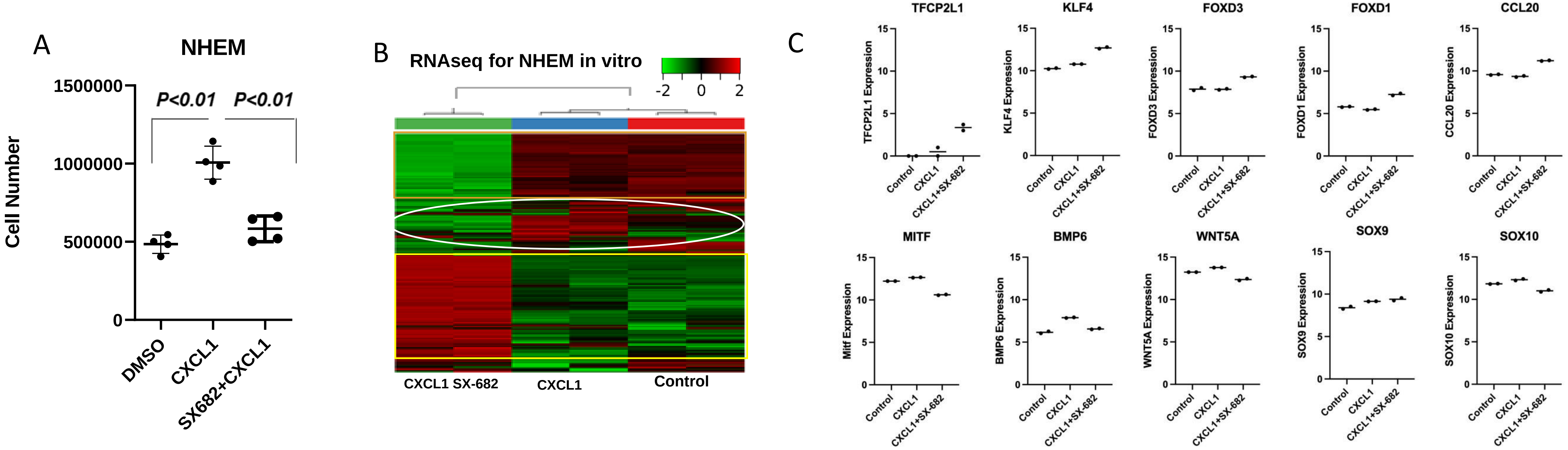
A
NHEM
Cell Number
1500000
1000000
500000
0
P<0.01
P<0.01
DMSO
CXCL1
SX682+CXCL1
B
RNAseq for NHEM in vitro
-2
0
2
CXCL1 SX-682
CXCL1
Control
C
TFCP2L1
KLF4
FOXD3
FOXD1
CCL20
MITF
BMP6
WNT5A
SOX9
SOX10
TFCP2L1 Expression
KLF4 Expression
FOXD3 Expression
FOXD1 Expression
CCL20 Expression
Mitf Expression
BMP6 Expression
WNT5A Expression
SOX9 Expression
SOX10 Expression
Control
CXCL1
CXCL1+SX-682

### Table S1

| Cytokines, Chemokines, and Interleukins | | |
|---|---|---|
| Gene | CXCL1 vs. Control log2FC (adj. p value) | CXCL1 + SX-682 vs. CXCL1 alone log2FC (adj. p value) |
| CCL18 | -0.388 (0.555) | -1.766 (9.67e-4) |
| CCL2 | -0.426 (1.19e-22) | -6.537 (<1e-310) |
| CCL20 | -0.298 (0.044) | 1.612 (1.42e-68) |
| CCL7 | -0.405 (0.052) | -9.397 (2.18e-10) |
| CCL8 | 0.009 (0.982) | -9.753 (4e-11) |
| CXCL1 | -0.345 (2.9e-19) | -4.033 (<1e-310) |
| CXCL10 | -1.476 (4.06e-9) | -1.471 (1.74e-4) |
| CXCL11 | -1.425 (2.3e-5) | -0.482 (0.324) |
| CXCL12 | 0.710 (3.74e-18) | -6.13 (1.05e-232) |
| CXCL2 | -0.002 (0.993) | -1.219 (2.24e-46) |
| CXCL3 | -0.361 (0.001) | -1.384 (2.45e-28) |
| CXCL5 | -0.62 (3.86e-36) | -1.604 (6.37e-181) |
| CXCL6 | -0.386 (5.38e-15) | -5.928 (<1e-310) |
| CXCL8 | -0.497 (7.99e-30) | -1.319 (5.89e-199) |
| IL11 | 1.001 (2.75e-23) | -0.285 (0.002) |
| IL11RA | 0.011 (0.97) | -1.622 (2.51e-21) |
| IL12RB2 | 0.51 (0.041) | -1.288 (2.57e-9) |
| IL13RA1 | -0.086 (0.233) | -0.358 (1.42e-12) |
| IL15RA | -0.37 (0.406) | 1.693 (1.46e-14) |
| IL16 | -0.267 (0.40) | -2.293 (1.01e-14) |
| IL17RB | 0.019 (0.971) | 0.739 (5.66e-4) |
| IL17RD | -0.190 (0.362) | -0.465 (0.003) |
| IL17RE | 0.229 (0.718) | -0.794 (0.042) |
| IL18BP | 0.052 (0.907) | 0.704 (3e-4) |
| IL18R1 | 0.884 (0.025) | 2.688 (4.89e-60) |
| IL1R1 | 0.293 (6.71e-11) | -2.017 (<1e-310) |
| IL1RL1 | 1.968 (5.41e-5) | 3.916 (6.28e-141) |
| IL24 | -0.123 (0.204) | -0.313 (1.38e-5) |
| IL27RA | -0.223 (0.316) | -0.601 (4.75e-4) |
| IL33 | 0.689 (0.037) | -4.154 (1.45e-21) |
| IL34 | -1.609 (7.45e-13) | -0.166 (0.639) |
| IL4R | 0.027 (0.791) | -0.494 (6.65e-22) |
| IL6 | 0.025 (0.771) | -0.82 (6.57e-77) |
| IL6R | 0.089 (0.59) | -1.611 (4.21e-51) |
| IL7 | -0.569 (0.023) | 1.122 (3.5e-10) |
| IL7R | -0.259 (0.487) | -1.025 (4.34e-4) |

### Table S2

| TNF Related Cytokines and Interferons | | |
|---|---|---|
| Gene | CXCL1 vs. Control log2FC (adj. p value) | CXCL1 + SX-682 vs. CXCL1 alone log2FC (adj. p value) |
| C1QTNF1 | -0.135 (0.008) | -1.446 (6.63e-243) |
| C1QTNF2 | -1.013 (0.017) | -4.923 (7.33e-8) |
| C1QTNF6 | 0.142 (0.246) | -2.543 (4.08e-117) |
| TNFAIP1 | 0.011 (0.921) | -0.22 (2.66e-5) |
| TNFAIP2 | -0.345 (0.004) | -1.283 (4.44e-24) |
| TNFAIP3 | -0.261 (5.54e-6) | -1.066 (1.15e-79) |
| TNFAIP6 | 0.044 (0.873) | -2.854 (6.99e-50) |
| TNFAIP8 | -0.707 (4.81e-18) | -1.323 (2.97e-38) |
| TNFAIP8L1 | -0.334 (0.044) | -2.102 (1.8e-33) |
| TNFAIP8L3 | -0.048 (0.696) | -0.894 (3.21e-36) |
| TNFRSF10A | 0.051 (0.822) | 0.307 (0.006) |
| TNFRSF10B | 0.012 (0.899) | 1.231 (4.85e-177) |
| TNFRSF10D | 0.223 (0.001) | -1.663 (2.06e-136) |
| TNFRSF11B | -0.168 (0.235) | -2.937 (8.15e-91) |
| TNFRSF12A | -0.194 (0.428) | 0.659 (8.34e-6) |
| TNFRSF14 | 0.245 (0.001) | 0.047 (0.515) |
| TNFRSF18 | -0.069 (0.942) | -2.208 (4.24e-4) |
| TNFRSF19 | 0.093 (0.344) | -2.891 (3.17e-230) |
| TNFRSF1A | 0.294 (2.96e-11) | -0.989 (6.79e-127) |
| TNFRSF1B | -0.151 (0.0507) | -1.757 (3.73e-131) |
| TNFRSF21 | 0.094 (0.10) | -3.004 (<1e-310) |
| TNFSF11 | 0.898 (1.46e-55) | -3.465 (<1e-310) |
| TNFSF12 | -0.315 (0.031) | -1.333 (2.23e-20) |
| TNFSF13B | 0.291 (0.145) | -3.421 (1.04e--53) |
| TNFSF4 | -0.493 (0.052) | 1.161 (2.51e-11) |
| IFNAR1 | -0.074 (0.346) | -0.785 (7.76e-49) |
| IFNAR2 | 0.132 (0.801) | -1.343 (2.31e-5) |
| IFNGR1 | 0.144 (0.026) | 0.059 (0.309) |
| IFNGR2 | -0.109 (0.193) | -0.612 (1.45e-22) |
| IFNLR1 | -0.022 (0.963) | -1.212 (4.26e-6) |

**Supplemental Figure Legends:**

Figure S1. Breakdown of CXCR2 (A) and CXCL1 (B) expression in nevi and melanoma from TCGA, separated based on BRAF and NRAS mutation status. Data were analyzed with a two-way ANOVA and no significant differences were found.

Figure S2. Diagram showing strategy for development of $Tyr\text{-}Cre^{ER}::Braf^{V600E}::Pten^{-/-}::mT/mG^{fl/fl}::Cxcr2^{fl/fl}$ and $Tyr\text{-}Cre^{ER+}::NRas^{Q61R}::Ink4a^{-/-}::mT/mG^{fl/fl}::Cxcr2^{fl/fl}$ mice (A, B, figures created using Biorender). Image of GFP-expressing $Braf^{V600E}/Pten^{-/-}$ tumors *in vivo* and via histology (C, D).

Figure S3. Heat maps of the most differentially expressed genes in tumors from the $Braf^{V600E}/Pten^{-/-}$ mice with or without loss of CXCR2 in tumor cells (A), with control or SX-682 treatment (B), and in MelanA, B16F0, and B16F10 cells treated with DMSO or SX-682 (C). Tumor suppressive genes are listed in red, genes involved in growth are in green, immune related genes are in blue, differentiation/stemness genes are in purple, and motility and cell adhesion genes are in brown.

Figure S4. Heat map showing genes involved in growth that are suppressed (A) and genes involved in growth suppression that are induced (B) in $Braf^{V600E}/Pten^{-/-}/Cxcr2^{-/-}$ melanoma tumors. Red arrows indicate genes involved in growth/oncogenes (A) and inhibition of growth/tumor suppression (B).

Figure S5. Additional data from analysis of immune cells and cytokines expressed by $Braf^{V600E}/Pten^{-/-}/Cxcr2^{-/-}$ and $Braf^{V600E}/Pten^{-/-}/Cxcr2^{WT}$ tumors. mMCPCounter predicted infiltrate of B-derived cells, memory B cells, neutrophils, endothelial cells, mast cells, basophils, eosinophils, and blood vessels (A). FACS analysis of peripheral blood CD45+

cells from $Braf^{V600E}/Pten^{-/-}/Cxcr2^{-/-}$ and $Braf^{V600E}/Pten^{-/-}/Cxcr2^{WT}$ mice (B). Results of cytokine array of tumor lysates from $Braf^{V600E}/Pten^{-/-}/Cxcr2^{-/-}$ and $Braf^{V600E}/Pten^{-/-}/Cxcr2^{WT}$ melanoma tumors, expressed as fold change in $Cxcr2^{-/-}$ compared to $Cxcr2^{WT}$(CCL20 removed and shown in Figure 3D to allow proper visualization) (C). FACS analysis of CD45+ cells from $Braf^{V600E}/Pten^{-/-}/Cxcr2^{-/-}$ and $Braf^{V600E}/Pten^{-/-}/Cxcr2^{WT}$ tumors (D). Statistical analyses: (A), (C), (D) Welch's t-test

Figure S6. Additional data from analysis of immune cells from SX-682-treated $Braf^{V600E}/Pten^{-/-}/Cxcr2^{WT}$ tumors. mMCPCounter predictions for T cells, B-derived cells, memory B cells, NK cells, endothelial, monocytes, macrophages, fibroblasts, lymphatics, mast, basophils, eosinophils, and blood vessels (A). FACS analysis of peripheral blood CD45+ cells from tumor-bearing mice fed SX-682 or control chow (B). FACS analysis of CD45+ cells in SX-682 treated or control tumors (C). Statistical analyses: (A), (B), (C) Welch's t-test

Figure S7. Effects of SX-682 on MelanA, B16F0, and B16F10 cell lines. MelanA, B16F0, and B16F10 cells were treated with SX-682 (5μM) or DMSO, and cell number was evaluated at days 1-4 post treatment. Data were analyzed on a natural log scale and compared using two-way ANOVA with BH correction for multiple tests (A). Cell lines were treated for 24 hours with SX-682 or DMSO prior to RNA isolation and sequencing. A heat map shows that genes were commonly suppressed or induced in all three cell lines (B). Reverse-phase phosphoproteome analysis (RPPA) was also performed on cells treated for 24 hours with SX-682 or DMSO. Volcano plot showing phosphoproteins commonly altered by SX-682 treatment in MelanA, B16F0, and B16F10 cells (C). A heat map showing differential expression of commonly suppressed or induced phosphoproteins in response to treatment with 5μM SX-682 for 24 hours (D). Cytokine array of cell lysates from MelanA, B16F0, and B16F10 cells treated with SX-682 (5μM) for 24 hours as compared to those treated with DMSO (E).

Figure S8. Expression of *Tfcp2l1*-related genes and tumor markers based on RNAseq analysis in *Braf*$^{V600E}$/*Pten*$^{-/-}$/*Cxcr2*$^{-/-}$ tumors, *Braf*$^{V600E}$/*Pten*$^{-/-}$ tumors treated with SX-682, compared to appropriate controls. Statistical analysis: Welch's t-test

Figure S9. Expression of Tfcp2l1-related genes based upon RT-PCR analysis. MelanA, B16F0, and B16F10 cells were treated with SX-682 (5μM) for 24 hours prior to RNA extraction, quantification, and RT-PCR using primers specific for *Tfcp2L1, Foxd3, Sox2, Sox10, Notch1, Hmga2, Mitf, Klf4, Myc, Nanog, Esrrb and Tyr*. Data are plotted as fold-change compared to DMSO treated cultures. Statistical analysis: Welch's t-test

Figure S10. WGCNA orders genes into a dendrogram by their co-expression profiles across the samples. Gene modules at the bottom of the dendrogram show assignment into distinct clusters of co-expressed genes (A). Module eigengene expression value shown as a heatmap across WT and KO samples (B). Feature selection of modules by ANOVA F-value (FDR-adjusted p-value). Six of 10 WGCNA gene modules can distinguish between conditions (C). Eigengene expression value across WT and KO samples shown as a bar-plot and significance of transcription factors (ordered by FDR-adjusted p-value) within each module for the six significant gene modules from panel D: blue, turquoise, brown, green, yellow, and red. These TFs are best at distinguishing between KO and WT samples in each module (D).

Figure S11. ChIPseq analysis of changes in gene promoter binding of TFCP2L1 antibody in B16F0 cells in response to SX-682 treatment. B16F0 cells were cultured overnight with either DMSO or SX-682. Cell extracts were incubated with anti-IgG or anti-TFCP2L1, cross-linked with a reversible cross-linker, and prepared and prepared for ChIP-seq analysis. Metascape analysis was used to examine enriched terms across input gene lists of SX-682 minus DMSO, colored by p-values. Top terms associated with TFCP2L1 bound promoters where gene expression was increased (A) and decreased (B) were identified. Metascape Analysis was used to compare the data sets from RPPA, ChIP, and RNAseq analyses of B16F0 cells treated with SX-682 as compared to respective controls (C). Cluster analysis of genes identified as regulated by SX-682 by ChIPseq, RNAseq, or RPPA, with commonly enriched GO terms across all three methods (D). Genes in Cluster analysis of S11D were colored by MCODE ID (E).

Figure S12. NHEM cells were cultured in melanocyte growth medium containing vehicle or CXCL1 (100ng/ml), or CXCL1 (100ng/ml) with 5 μM SX-682. After 5 days of culture, the cell number was determined, and total RNA was extracted for RNAseq analysis. Effects of SX-682 on cell growth in vitro (A). Heatmap for RNAseq on NHEM. Five days after NHEMs were cultured with SX-682 and/or CXCL1, RNA was extracted and subjected to RNAseq analysis(B). Expression values for key genes associated with response to CXCL1 or CXCL1 and SX-682 (C).

Table S1. Comparison of Cytokine, Chemokine, and Interleukin Expression Following CXCL1 or CXCL1+SX-682 Treatment. Key cytokines induced are highlighted in red and reduced are highlighted in green.

Table S2. Comparison of TNF-Related Cytokines and Interferon Expression Following CXCL1 or CXCL1+SX-682 Treatment. Key cytokines induced are highlighted in red and reduced are highlighted in green.
